# Systemic coordination of whole-body tissue remodeling in sea anemone local regeneration

**DOI:** 10.1101/2024.03.20.585927

**Authors:** Stephanie Cheung, Danila Bredikhin, Tobias Gerber, Petrus J. Steenbergen, Soham Basu, Richard Bailleul, Pauline Hansen, Alexandre Paix, Matthew A. Benton, Hendrik C. Korswagen, Detlev Arendt, Oliver Stegle, Aissam Ikmi

**Affiliations:** European Molecular Biology Laboratory (EMBL), Developmental Biology Unit, Heidelberg, Germany; European Molecular Biology Laboratory (EMBL), Genome Biology Unit, Heidelberg, Germany; Division of Computational Genomics and Systems Genetics, German Cancer Research Center (DKFZ), Heidelberg, Germany; Hubrecht Institute, Royal Netherlands Academy of Arts and Sciences and University Medical Center Utrecht, Utrecht, Netherlands; Faculty of Biosciences, Heidelberg University, Heidelberg, Germany; Wellcome Sanger Institute, Wellcome Genome Campus, Hinxton, United Kingdom

## Abstract

The complexity of regeneration extends beyond local wound responses (1,2), eliciting systemic processes that engage the entire organism. However, their functional relevance, and the spatial and temporal orchestration of the underlying molecular processes distant from the injury site remain unknown. Here, we demonstrate that local regeneration in the cnidarian *Nematostella vectensis* involves a systemic homeostatic response, leading to coordinated whole-body remodeling. Leveraging spatial transcriptomics, endogenous protein tagging, and live imaging, we comprehensively dissect this systemic response at the organismal scale. We identify proteolysis as a critical process driven by both local and systemic upregulation of metalloproteases. We show that metalloproteinase expression levels and activity scale with the extent of tissue loss, leading to proportional long-range movement of tissue and its associated extracellular matrix. Our findings illuminate the adaptive nature of the systematic response in regeneration. We propose that this integrated regenerative mechanism, shifting the system from a steady to a dynamic homeostatic state, allows the organism to cope with a wide range of injuries.

## Introduction

The regenerative capacity in the animal kingdom is remarkably diverse (3,4), encompassing extensive whole-body regeneration in invertebrates such as cnidarians and planarians, in stark contrast to the more restricted, tissue-specific regeneration in vertebrates (5-7) (Fig. 1a). This array of regenerative phenomena reflects a broad spectrum of biological resilience across animal lineages. Despite this diversity, a common thread is the initiation of both localized and systemic responses to tissue injury, manifested through a series of cellular and molecular events (8-12).

**Fig. 1.**
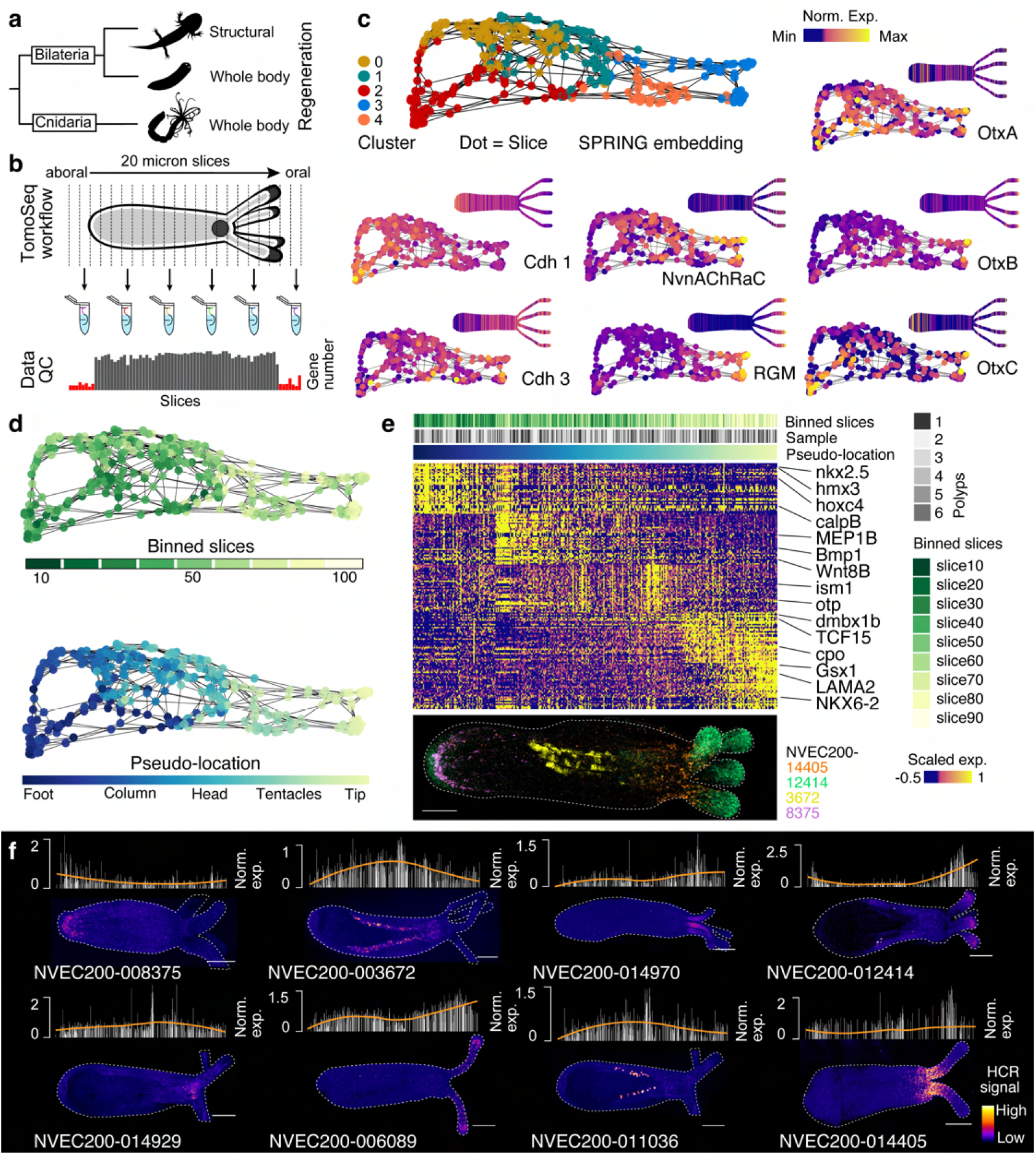
A transcriptome-based coordinate system along the oral-aboral axis of intact polyps. (**a**) A simplified phylogenetic tree showing the placement of Cnidaria as the sister group to Bilateria and the regeneration capacity of selected species, including a salamander and a planarian. (**b**) Schematic summarizing the tomo-seq experimental workflow. (**c**) SPRING embedding of 331 tomo-seq slices from six intact, primary polyps (top left). Color denotes Louvain cluster identities. Grey lines show the connection of slice transcriptomes in the SPRING k-nearest-neighbor graph. Insets: Expression levels of known axial gene markers are shown as features on top of the embedding and as in silico polyps (see also Fig. S2). (**d**) Top: Binned slice indices are visualized on top of the embedding. 10 slices are united per bin. Bottom: A pseudo-location estimated using a diffusion map algorithm visualized on top of the SPRING embedding. (**e**) Top: Heatmap representation of gene expression values for top 30 cluster-specific markers. Slices are ordered by pseudo-location presented in d. Bottom: Clusters mark different body regions as validated by a maximum projection image of whole-mount HCR *in situ* for 4 identified marker genes. Scale bar: 100 µm. (**f**) Maximum projection images of whole mount HCR *in situ* for 8 marker genes expressed in different anatomical regions. Their corresponding expression patterns in the tomo-seq data are shown as histograms above each HCR image. Slices (x-axis) in the histograms are ordered by pseudo-location, and a loess fitted line resembles the expected HCR pattern (see also Figure S2). HCR intensity profiles are shown as pseudocolor palette ‘fire’. Scale bars: 100 µm.

In regenerative models such as the Axolotl, limb amputation triggers cell cycle reactivation not only at the wound site but also in distant organs, including the contralateral limb, spinal cord, liver, and heart (11). Analogous phenomena are observed in mice, where injury activates quiescent stem cells in remote tissues (10), and in zebrafish, where cardiac injury induces gene expression changes in distant organs like the brain (9). Invertebrates also display similar patterns, with planarians and acoels showing initial body-wide cellular proliferation post-injury that becomes concentrated at the regeneration site (8,13). Moreover, interfering with long-range signaling can affect regenerative outcomes, as observed in planarians (14). Such insights highlight the need for a holistic approach to understanding the complex processes underlying regeneration across different animal taxa.

Our study focuses on the starlet sea anemone, *Nematostella vectensis*, a cnidarian known for its extensive whole-body regeneration capabilities (15,16). We leverage its genetic tractability (17,18) to conduct an in-depth investigation of regeneration, comparing intact and regenerating animals, from genome-wide expression patterns and molecular activities to tissue dynamics and overall organismal outcomes. The relatively simple morphology yet complex tissue architecture of the *Nematostella* polyp, with its bi-layered tissue and interstitial mesoglea (also known as extracellular matrix, ECM), supports a diverse range of cell types along its body axis (19,20). Key cellular mechanisms, particularly proliferation and apoptosis, play critical roles in oral regeneration (15,16,21).

## Results

### Spatial transcriptomics of intact polyps

To characterize spatial gene expression patterns in intact animals, we employed tomo-seq on primary polyps along the oral-aboral axis, each measuring around 1 mm in length (Fig. 1b and Fig. S1a). Briefly, this technique combines tissue cryo-sectioning with RNA sequencing on individual sections to generate an expression atlas with uni-dimensional spatial information (22-24). We processed six polyps at a resolution of 20 µm slices, yielding a total of 331 slices (Fig. 1b; Fig. S1a and Table 1; following quality control, yielding 50-60 slices per polyp). To accommodate variations in animal length and resulting slice numbers, we considered two complementary strategies for aligning the spatial transcriptomic data across polyps.

We first conducted reference-based alignment based on dynamic time warping (25) to align slices from all polyps to a common coordinate system defined based on a pre-selected high-quality reference polyp (Fig. S1b-c). Anatomical marker genes identified as spatially variable in the reference polyp showed consistent spatial expression patterns across the aligned polyp slices, delineating spatial domains along the oral-aboral axis (Fig. S1d-e and Table 2). As a second strategy, with the aim to avoid potential bias towards a single designated reference polyp, we used Harmony (26) to jointly integrate the transcriptomes from all polyp slices, followed by clustering of slices based on the aligned harmony components (Fig. S1f). Visualization of these clusters on a SPRING embedding (27) revealed five distinct gene expression states (Fig. 1c-e and Fig. S1f). We next used a diffusion map approach to estimate transcriptome-based pseudo-locations for each slice across all polyps (28) (Fig. 1f). Remarkably, the pseudo-ordering broadly reflected the original slice ordering (Fig. S1g,h), with minor differences occurring for lower-quality slices (Fig. S1i). Moreover, mapping previously reported genes’ expression patterns onto the embedding also supported that the pseudo-ordering aligns with known oral-aboral markers’ expression patterns, such as Cadherin 1/3 (29), Otx (30), RGM (31) and nAChRc (32) (Fig. 1c), validating our approach. We also used the reference-free, pseudoaligned data to identify additional genes that vary across the body (Wilcoxon Rank Sum test, log2FC > 0.3; Fig. 1e, Fig. S1j and Table 2).

To validate these findings, we performed an *in situ* hybridization chain reaction (HCR) for several genes showing distinct spatial expression patterns across the polyps (Fig. 1e,f). Out of 16 genes tested, 15 displayed the expected expression pattern (Fig. 1f and Fig. S2), with the exception of one gene (NVEC200-006163: HSPA12A) for which no *in situ* signal was detected. Collectively, these findings demonstrate that the spatially aligned transcriptomic maps effectively identify robust gene expression domains along the oral-aboral axis of primary polyps.

### Spatiotemporal transcriptomics of regenerating polyps

Leveraging spatial transcriptomic maps created for intact animals, we sought to understand whether dynamic changes in gene expression occur throughout the entire body of regenerating polyps. We conducted surgical foot removal, cutting approximately 20% of the polyp, followed by transcriptomic investigation, and tracked the reestablishment of Fgfrb-eGFP expressing cells (indicators of the aboral pole tip) as a marker of complete foot regeneration (Fig. 2a). Fgfrb-eGFP expressing cells began to appear around 48 hours post-amputation (hpa) and were fully restored at 96 hpa. We collected and analyzed slices from three to four polyps per time point at 12, 24, 48, and 96 hpa (Fig. S3a and Table 1, following quality control, yielding 36-56 slices per polyp).

**Fig. 2.**
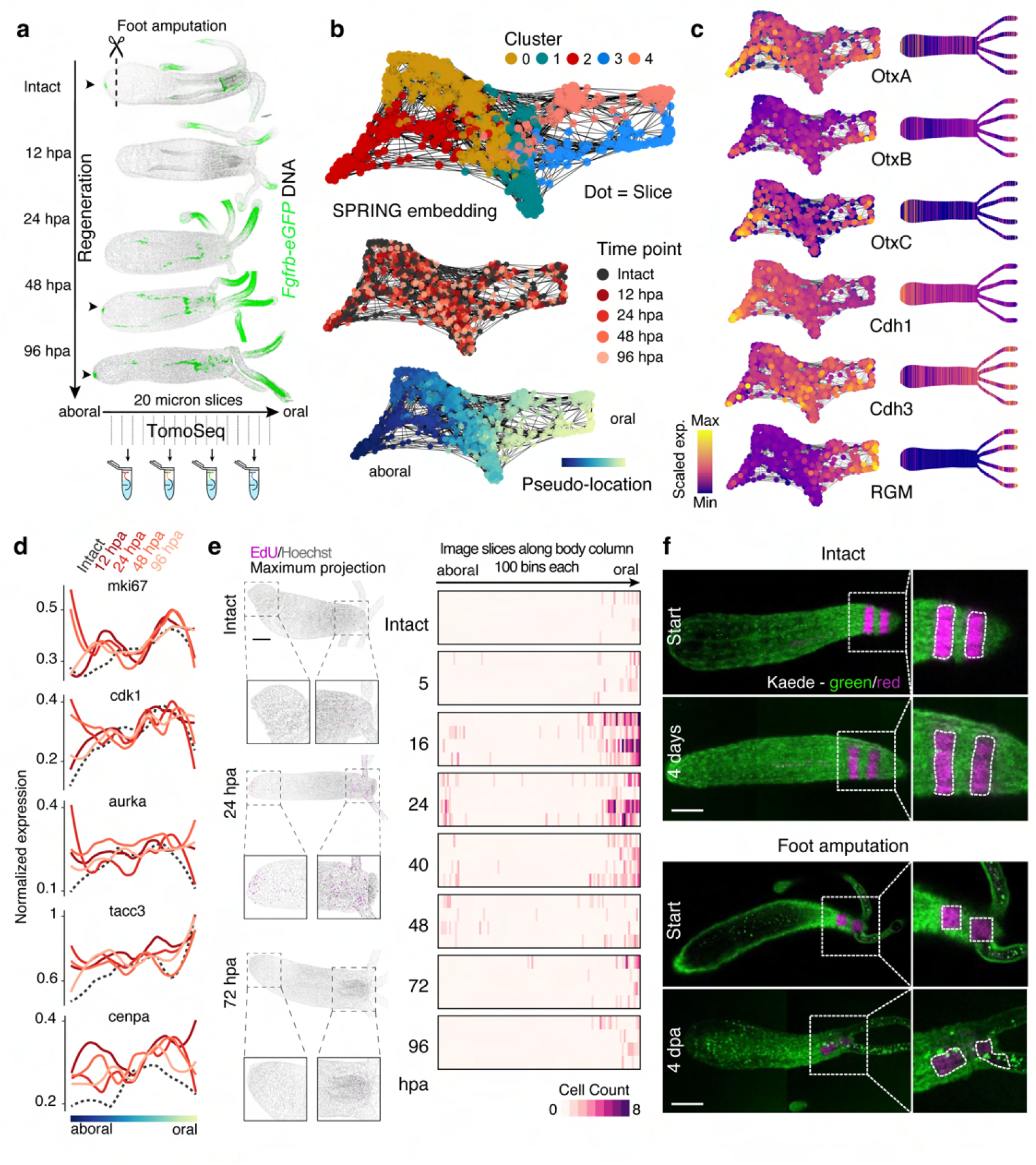
Mapping spatiotemporal transcriptomic changes and tissue dynamics in foot-regenerating polyps. (**a**) Maximum projection images of transgenic Fgfrb-eGFP polyps at indicated time points following foot amputation (top) and a schematic of the tomo-seq workflow (bottom). Green: Fgfrb-eGFP. Gray: DNA. Scale bar = 50 microns. (**b**) Top: SPRING embedding of 957 tomo-seq slices of intact and regenerating primary polyps (intact, n= 6 polyps; 12hpa, n=4 polyps; 24hpa, n= 3 polyps; 48hpa, n= 3 polyps; 96hpa, n= 4 polyps). Colors represent Louvian cluster identity. Grey lines show the connection of slice transcriptomes in the SPRING k-nearest-neighbor graph. Center: All time points are visualized on top of the embedding. Bottom: A pseudo-location ranking calculated by a diffusion map algorithm is visualized on top of the embedding. (**c**) Expression levels of known axial gene markers are shown as features on the embedding (left) and as in silico polyps (right). (**d**) Expression of cell cycle marker genes is visualized as a smoothed curve across pseudo-ordered slices for each time point. (**e**) Left: Maximum projection images of representative polyps showing EdU-positive cells (magenta) and nuclei (grey) at the indicated time points. Inset boxes are zoom-in views of the oral and aboral poles. Right: Heatmaps visualize the number of EdU-positive cells at different time points per bin (columns) for 4 polyps (rows) for each time point. 100 bins were determined for each polyp along the aboral-oral axis excluding the tentacles (see Fig. S4). (**f**) Maximum projection images of photo-converted oral tissue patches in intact (top) and foot-regenerating (bottom) polyps expressing Kaede (green: not converted; magenta: photo-converted). The pattern of the photo-converted patches is highlighted in dashed lines and shown shortly after the photoconversion (start) and four days post-conversion. Inset boxes are zoom-in views of the oral pole. Scale bars: 100 µm.

To study local and global effects of surgical foot removal, we jointly integrated the tomo-seq slices from intact and regenerating animals using the reference-free Harmony approach, followed by clustering and pseud-time inference (Fig. 2b and Fig. S3b-c), giving rise to a dataset with 957 integrated slices. Consistent with the results from intact animals, the aboral-oral axis was identified as the main source of gene expression variation among tomo-seq slices also in regenerating animals (Fig. 2b and Fig. S3c). The joint embedding generated a coherent coordinate system across intact and amputated animals (Fig. 2b and Fig. S3d-g). The persistence of axial marker expression in regenerating polyps further substantiated this observation (Fig. 2c and Fig. S3h-j). Thus, we generated robust spatiotemporal molecular maps for regenerating animals.

### Dual orchestration of morphallactic and epimorphic processes

As the onset of cell proliferation often demarcates the transition from wound healing to regeneration, we initially considered spatial expression maps of known cell cycle genes (Fig. 2d and Fig. S4a,b). As expected, canonical cell cycle markers such as MKI67, CDK1, AURKA, TACC3, and CENPA showed upregulation in the regenerating tissue at 12 and 24 hpa Fig. 2d), indicating the involvement of cell proliferation during foot regeneration (33). However, a particularly intriguing observation was the simultaneous gradual upregulation of these genes towards the oral pole, suggesting an additional distal response to tissue damage even though the oral tissue was not damaged (Fig. 2d and Fig. S4b). To confirm this proliferation pattern, we employed EdU labeling, marking cells in the S-phase along the oral-aboral axis within a timeframe of 5 to 96 hpa (Fig. 2e and Fig. S4c,d). Consistent with the tomo-seq results, we observed an increase in EdU+ cells at the same time points at the wound site (12 and 16 hpa; Fig. 2e and Fig. S4d). Moreover, we observed a marked increase in EdU+ cells towards the oral pole, again confirming the observations from spatial transcriptomics, reinforcing the notion of proliferation being stimulated in regions distant from the immediate site of regeneration.

Motivated by the observation in the cnidarian *Hydractinia* that remote proliferative cells migrate toward the regenerating tissue (34), we performed a photoconversion experiment to monitor the movement of photo-converted Kaede-positive cells at the oral pole following amputation. (Fig. 2f and Fig. S4e,f). No discernible movement of photo-converted oral cells was observed toward the regenerating foot. Instead, photo-converted patches of cells reorganized along the main axis, and in some cases, patches in near proximity to the oral pole moved towards the tentacles (Fig. 2f and Fig. S4e,f), reflecting orientated cell rearrangement. One prediction of such morphallactic remodeling is the readjustment of the aspect ratio of the body, which is consistent with observed results (Fig. S5). In light of these findings, the activation of both wound localized and distal cell proliferation, coupled with tissue remodeling, illustrates a complex systemic regenerative mechanism, indicative of a dynamic shift in homeostatic balance.

### Molecular landscape of systemic homeostatic response

Having established the dual deployment of morphallactic and epimorphic processes during foot regeneration, we next set out to identify molecular processes that change in response to amputation (Fig. 3a). We conducted a global body-wide differential expression analysis, separately considering an early regenerative stage (12hpa/24hpa), a mid-regenerative stage (24hpa/48hpa) and a late stage (48hpa/96hpa). To identify stage-specific molecular processes, we then tested for positively differentially expressed genes (DE) at each stage versus all other time points as well as the intact animal (Early: N=215, Mid: N=216, Late: N=189,Wilcoxon Rank Sum test, log2FC > 0.5; Table 3). Gene Ontology (GO) enrichment analysis (top 50 genes per stage; GOseq p value < 0.05; Fig. 3c; Fig. S6a and Table 3) identified distinct molecular processes that mark these three regeneration stages (Fig. 3b,c). The early phase of regeneration was marked by an increase in proteolytic genes, including metallopeptidase ADAMTS6 and BP10 (Fig. 3b,d). This observation is consistent with previous transcriptomic studies on wound or regenerating polyps (12,35), highlighting their potential role in remodeling the ECM and processing growth factors. Transitioning to the mid-regenerative phase, a second set of genes became prominent, notably involved in cell communication, exemplified by Fras1 and Frem2. These genes are recognized for their contributions to basement membrane integrity in other systems (36,37). Finally, the late regeneration was characterized by an upregulation of G protein-coupled receptors, including Gamma-aminobutyric acid B receptor 2 (GABBR2), and Adhesion G Protein-Coupled Receptor V1 (ADGRV1). The sequential activation of distinct molecular events across regeneration stages demonstrates a coordinated molecular cascade of the systemic response in regeneration.

**Fig. 3.**
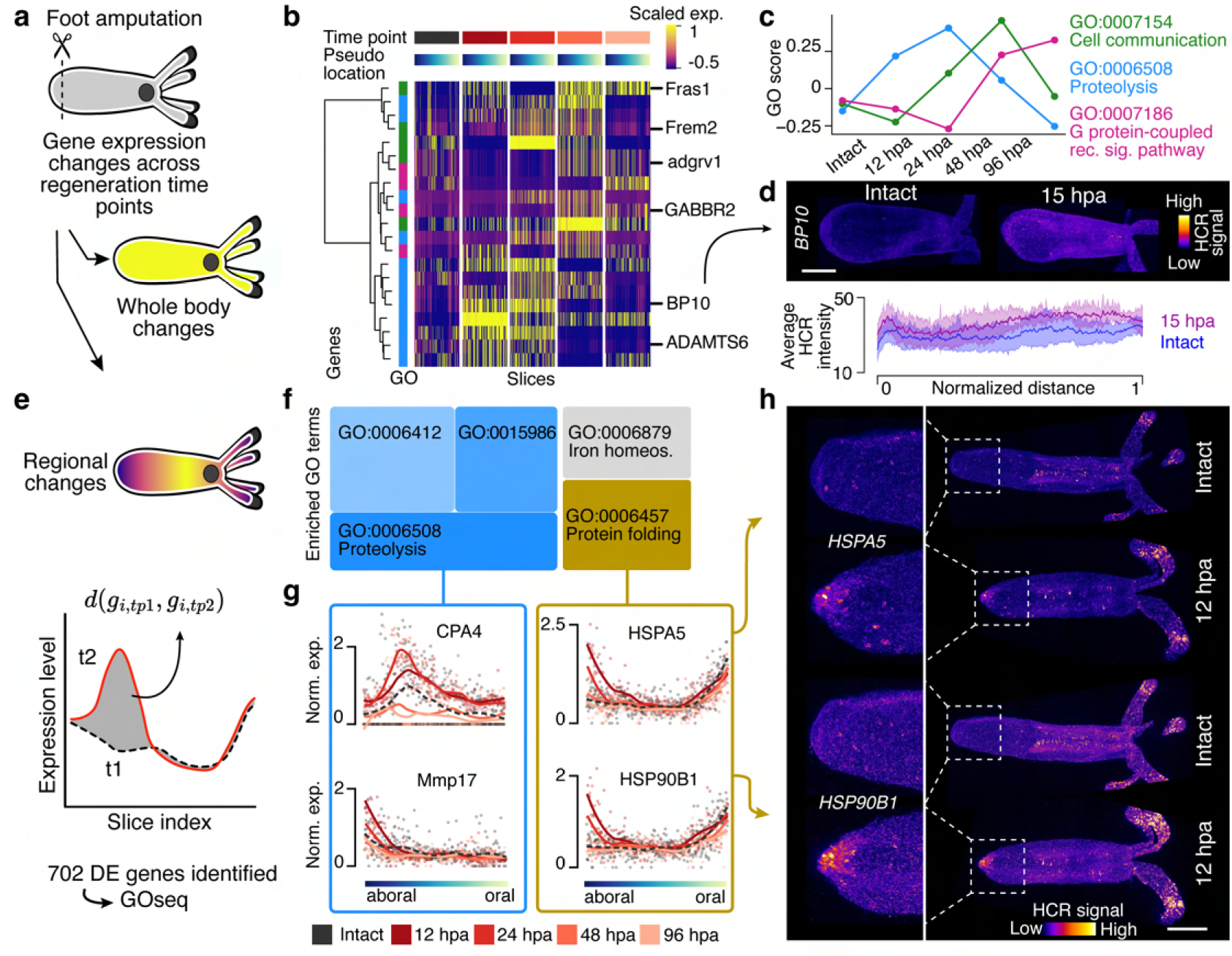
Proteolysis correlates with local and systemic wound responses. (**a**) Schematic illustration of alternative analysis strategies of gene expression changes during foot regeneration, either considering a global (body-wide) scale (panels b-d) or a local scale (panels e-h). (**b**) Heatmap showing relative gene expression levels for selected genes with evidence for global differential expression associated with different GO terms as in c. Genes correspond to rows; columns denote slice index for different time points. Slices are separated by time point and ordered by pseudo-location ranking (see Fig. 2b). Hierarchical clustering of genes across all slices shows broad separation of GO terms. Blue: GO:0006508, Green: GO:0007154, Magenta: GO:0007186. (**c**) Lineplot shows the progression of GO scores across time points. Scores are calculated based on genes in b. (**d**) Top: Maximum projection images of whole mount HCR *in situ* for the protease BP10 in intact and foot-regenerating polyps (15 dpa). Scale bar: 100 µm. Bottom: quantification of BP10 expression is shown as average intensity against the normalized distance. The shaded regions around each line represented the standard deviation around the mean (intact, n= 8 polyps; 15hpa, n= 7 polyps). (**e**) Schematic illustrating the analysis of spatially variable gene expression changes (spatial DE genes) in response induced by foot cut using a Gaussian process regression model. (**f**) GO enrichment analysis of the 702 spatial DE genes (using GOseq (39)). Shades of the same color combine terms that belong to a similar term category. (**g**) Scatter plot and interpolation of gene expression as a function of spatial location for different time points. Solid lines denote fitted loess curve fits. (**h**) Maximum projection images of whole mount HCR *in situ* for two heat shock proteins in intact and foot amputated animals at 12 hpa. HCR intensity profiles are shown as pseudocolor palette ‘fire’. Scale bar: 100 µm.

We further assessed whether gene expression changes in response to the injury also occurred in more localized patterns across the body. To investigate the spatial dynamics of gene expression during regeneration, we adapted a Gaussian process model (38) to test for positionally local differential expressed genes (spatial DE genes) between each regeneration time point and the intact reference (Fig. 3e). Across all time points, this analysis identified 702 spatial DE genes (FDR<1%) (Fig. S6b). Remarkably, 35% of these genes had evidence for differential spatial expression across all regeneration stages (Fig. S6b). GO enrichment analysis (39) identified Proteolysis (p-value < 0.02; Fig. 3f; Fig. S6c), which is in common with the global response process, but points also to Protein folding, and Iron homeostasis, which were exclusively identified by the local differential expression analysis.

Within these categories, diverse patterns of spatial DE were identified (Fig. S6c). For example, the carboxypeptidases CAP2 and CPA4 were differentially expressed away from the wound site while the matrix metalloproteinase 17 (Mmp17) was spatially differentially expressed at the wound site, and not expressed elsewhere. Other genes, such as HSPA5 and HSP90B1 also showed evidence for localized DE at the wound site, although these genes are also expressed in the oral pole in intact animals (Fig. 3g,h). Beyond the recognized stress response role of heat shock genes in regeneration (40-43), their additional expression patterns hint at a broader physiological significance in polyp biology. These findings underscore a complex molecular landscape of the systemic response in regeneration, shifting the organism from a steady to a dynamic state, with Proteolysis involved in diverse spatial response patterns.

### Local and systemic regulation of metalloproteases

Upregulation of proteolysis genes, as identified in both the global and local analysis of the gene expression response to injury, led us to hypothesize a widespread activation of metalloproteases across the organism. To test this hypothesis, we employed *in vivo* zymography (44), a method that allows for the spatial visualization of proteolytic activity through the cleavage of a dye-quenched (DQ-) substrate, resulting in fluorescence (Fig. 4a,b and Fig. S7a). At 15 hours post incubation with the DQ-Col4, intact polyps exhibited a basal level of fluorescence signal, indicative of normal homeostatic metalloprotease activities. In contrast, polyps subjected to foot amputation demonstrated a widespread and intense fluorescence signal distribution, suggesting a systemic induction of hydrolytic enzyme activities in regeneration, despite the localized nature of the injury. Intriguingly, this upregulation was even more enhanced in response to an increase in tissue loss following mid-body amputation (Fig. 4a,b and Fig. S7a), reflecting the extensive tissue remodeling demands following more severe damage.

**Fig. 4.**
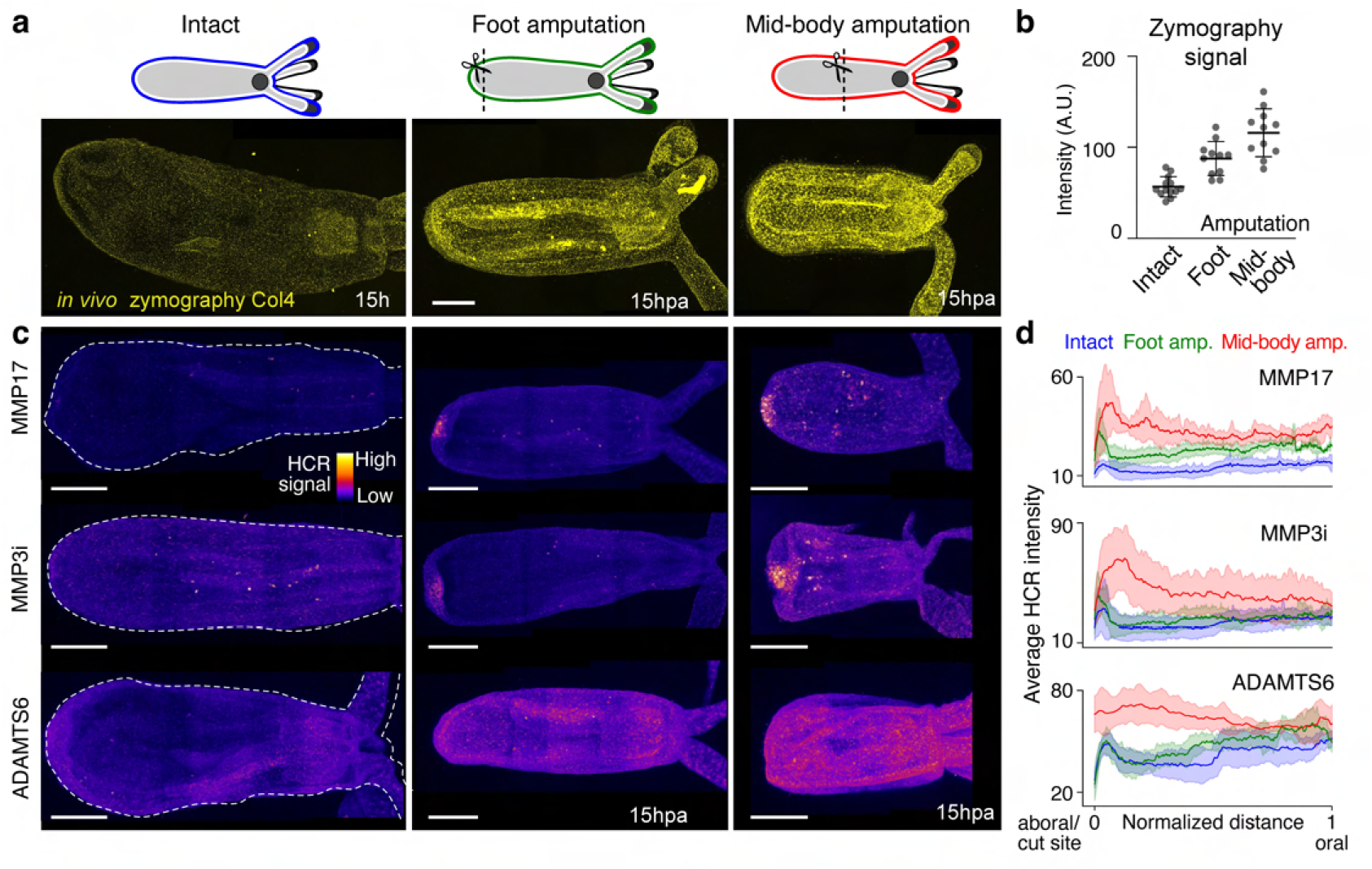
Local and systemic regulation of metalloproteases in regeneration. (**a**) Maximum projection images of intact, foot and mid-body amputated polyps incubated with DQ-Collagen 4 for *in vivo* zymography. Scale bar: 100 µm. (**b**) Quantification of de-quenched Collagen IV signal in the indicated conditions (intact, n= 13 polyps; foot amputation, n= 11 polyps; mid-body amputation, n= 11 polyps). Data are mean with SD for error bars. (**c**) Maximum projection images of whole mount HCR *in situ* for MMP17, MMP3i and ADAMTS6 in intact (left), foot amputated (center), and mid-body amputated polyps at 15hpa (right). See Fig. S7 for more representative examples. HCR intensity profiles are shown as pseudocolor palette ‘fire’. Scale bar: 100 µm. (**d**) Average expression levels of specified genes (MMP17, MMP3i, ADAMTS6) are plotted against normalized distances for each experimental condition: intact (blue), foot amputation (green), and mid-body amputation (red). The shaded regions around each line denote the standard deviation. Note that MMP17 expression in foot amputation also shows a slight systemic upregulation that was not detected in the tomo-seq data. Sample sizes are as follows: for MMP17, n= 7 intact polyps, n= 7 foot amputated polyps, n= 9 mid-body amputated polyps; for MMP3i, n= 6 intact polyps, n= 8 foot amputated polyps, n= 10 mid-body amputated polyps; for ADAMTS6, n= 8 intact polyps, n= 8 foot amputated polyps, n= 9 mid-body amputated polyps.

To link the changes in proteolytic activity levels to the identified gene expression response, we analyzed the expression patterns of both genes with local response at the wound site as well as systematic body-responding metalloenzymes using HCR (Fig. 4c). Consistent with the tomo-seq data, the expression of Mmp17 and its associated Matrix metalloprotease inhibitor 3 (Mmpi3) was activated at the wound site, whereas ADAMTS6 was ubiquitously upregulated in response to foot amputation (Fig. 4c,d and Fig. S7b-d). More strikingly, following mid-body amputation, not only did the expression levels of Mmp17 and Mmpi3 intensify at the wound site, but also triggered an additional systemic increase throughout the organism, including the tentacles (Fig. 4d). A similar trend was observed in the systemic expression of ADAMTS6, which again exhibited a more pronounced systematic response compared to the case of foot amputation (Fig. 4d). These findings demonstrate a coordinated, scaled response in metalloprotease expression and activity, aligning with the extent of tissue loss and highlighting the adaptive nature of systemic regulatory mechanisms in regeneration.

### Systemic remodeling of the basement membrane

To elucidate the downstream effects of metalloprotease activity on regeneration, it is crucial to study the ECM, particularly since metalloproteases are known to modulate ECM components. To assess the influence of metalloprotease activities on ECM protein dynamics, we developed a genetically modified line where Collagen 4 (Col4) is endogenously tagged with Dendra2 using CRISPR/Cas9 (Fig. 5a) (18). This modification specifically labels the basement membrane (18), enabling the *in vivo* observation of Col4::Dendra2 dynamics after photoconversion (Fig. 5a,b). We photoconverted eleven polyps, creating four spatially distinct photoconverted Col4::Dendra2 patches along the oral-aboral axis in each animal. We subjected six polys to foot amputation and monitored changes in the spatial position of the photoconverted patches across regeneration. Comparison of these amputated animals to intact animals (Fig. 5c and Fig. S8a) enabled us to estimate the velocity of displacement of these patches in response to injury (Fig. 5b,c). By 18 hours post-amputation, there was a significant global displacement of the Col4::Dendra2 patches in foot-amputated animals (Fig. 5c), in contrast to the minimal displacement observed in intact polyps. Notably, in regenerating animals, the photoconverted Col4::Dendra2 patches exhibited bidirectional movement: patches near the mouth moved orally, whereas those located lower on the polyp moved towards the aboral pole. This movement was also associated with a decrease in the intensity profile of the patches (Fig. 5c). These results indicate that the systemic activation of metalloproteases post-amputation leads to extensive remodeling of the basement membrane. Furthermore, this global change in ECM dynamics mirrored distal epithelial rearrangement (Fig. 2f and Fig. S4e,f), implying that during foot regeneration, both epithelial cells and the ECM are displaced in a coordinated manner, functioning as a composite structure.

**Fig. 5.**
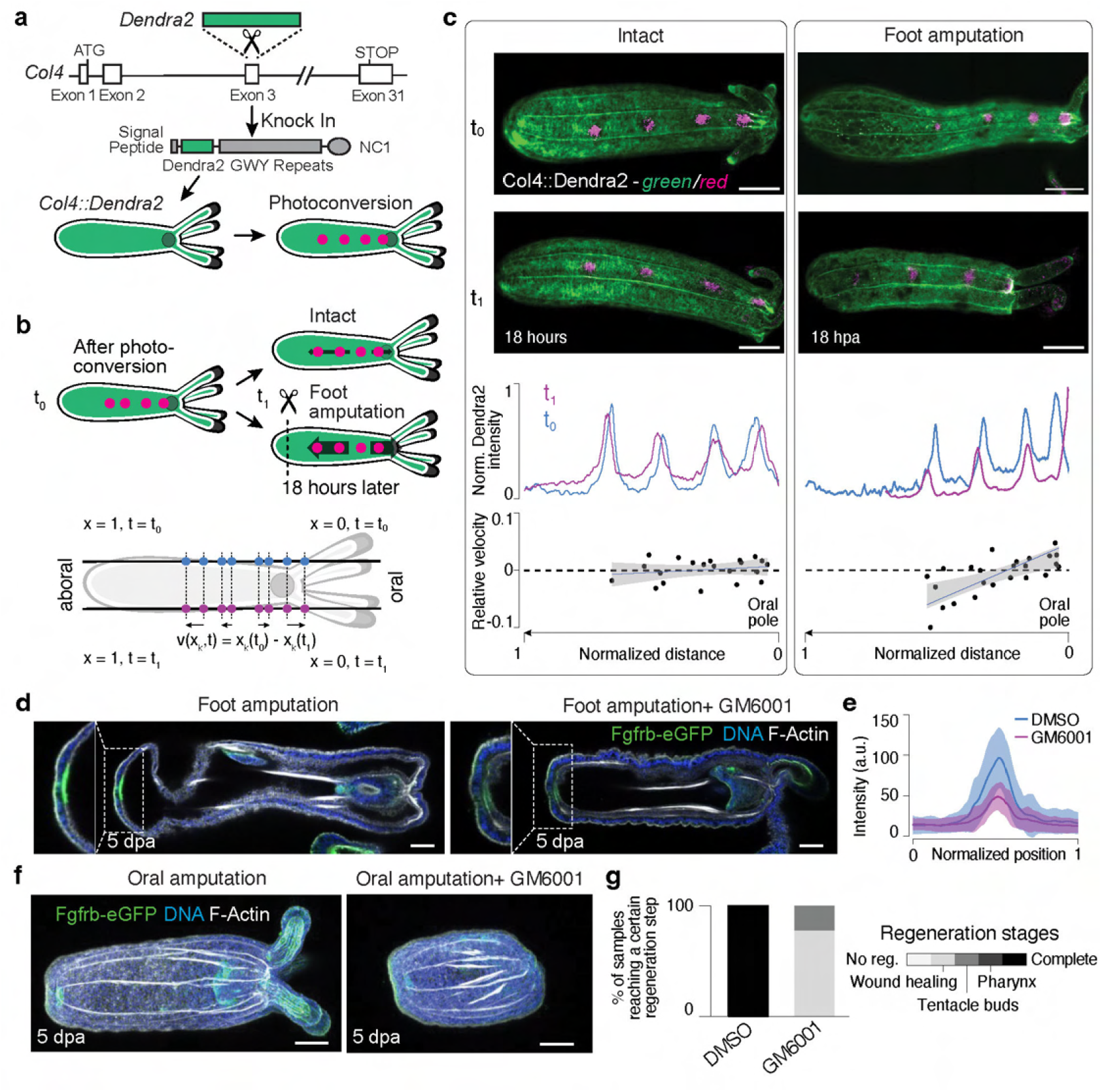
Local regeneration elicits systemic ECM remodeling. (**a**) Endogenous tagging of Col4 with the CRISPR/Cas 9 system. Dendra2 was inserted after the signal peptide which allows a precise site-specific photoconversion from green to red. The structural motifs of Col4 are shown. (**b**) Top: The experimental workflow of the photoconversion followed by a foot amputation with the Col4::Dendra2 polyps. Bottom: Illustration of the approach to estimate the velocity of patch movements along the oral-aboral axis in intact and regenerating animals. (**c**) Top: Maximum projection images of polyps showing Col4::Dendra2 (green) and photoconverted Col4::Dendra2 patches (magenta) along the oral-aboral axis in intact (left) and foot regenerating animals (right). t0: 0 hour, t1: 18 hours. Scale bars: 100 µm. See Fig. S8 for more representative examples. Bottom: Quantification of patch distances from the oral pole in intact (left, n= 5 polyps) and foot regenerating polyps (right, n= 6 polyps) at the indicated time points (top panels). Relative velocity estimations in intact (left) and regenerating animals (right) (bottom panels). 0 = Oral pole. 1 = Maximum normalized distance from the oral pole. (**d**) Cross-section images of a control polyp (left) and a GM6001-treated polyp (right) showing the expression pattern of Fgfrb-eGFP at the aboral tip (inset box) after foot amputation at 5 dpa. Scale bar: 50 µm. (**e**) Quantification of Fgfrb-eGFP expression. Line plot shows reduced GFP intensity after GM6001 treatment (DMSO control, n= 19 polyps; GM6001, n= 20 polyps). (**f**) Maximum projection images of a control polyp (left) and a GM6001-treated polyp (right) after oral amputation at 5 dpa. Scale bar: 50 µm. (**g**) Quantification of regeneration success in the indicated conditions (DMSO control, n= 8 polyps; GM6001, n= 11 polyps).

To explore the role of ECM remodeling in regeneration, we treated both foot-amputated and orally-amputated Fgfrb-eGFP polyps with a matrix metalloprotease (MMP) inhibitor, GM6001. Following treatment, wound closure was observed in both scenarios, effectively restoring epithelial integrity (Fig. 5d-g and Fig. S8c,d). However, during foot regeneration, Fgfrb-eGFP expression levels in GM6001-treated polyps were only partially restored compared to controls (Fig. 5e). Moreover, oral regeneration in polyps treated with GM6001 predominantly arrested at the wound-healing stage (Fig. 5f,g), with only a minority advancing to the early bud stage. These findings show the pivotal role of matrix metalloproteases in ECM remodeling and their significant impact on the morphogenetic processes essential for regeneration.

## Discussion

The field of regeneration has traditionally focused on wound-localized processes. Here, we provide a mechanistic understanding of how regeneration extends far beyond the wound site, involving a cascade of systemic processes that orchestrate coordinated whole-body remodeling. To elucidate these intricate mechanisms, we combined spatial transcriptomics, endogenous protein tagging, and live imaging, comparing intact and foot-regenerating polyps. These data serve as invaluable resources to gain deeper insights into the inherent biology of a highly regenerative species and offer new perspectives on the complex mechanisms underpinning regeneration.

A central finding of our study highlights the significant role of proteolysis, driven by local and systemic upregulation of metalloproteinases in regeneration. The expression levels and activity of these enzymes scale with the extent of tissue loss, enabling an adaptive, coordinated remodeling of the whole animal in regeneration. This proportionate effect was also reflected in the extent of bidirectional flow of both epithelial cells and their underlying ECM. This bidirectional pattern mirrors similar dynamics in Hydra, where continuous tissue and mesoglea movements are part of homeostasis (45,46). In *Nematostella*, however, such movement is triggered by injury, showing the dynamic nature of the cnidarian mesoglea. Previous work has identified localized ECM changes in the regeneration of other organisms (47-50), but our study brings attention to body-wide ECM changes governed by metalloproteases and leading to the gradual reshaping of the whole-body after tissue loss. We propose that this mechanism, pivotal in facilitating extensive remodeling, is likely a common strategy in species capable of whole-body regeneration.

Our work also uncovered a surprising induction of local cell proliferation at the oral pole in response to foot amputation, despite these cells not contributing directly to the regenerating tissue. This observation, coupled with the identification of diverse spatiotemporal gene expression responses and the long-range tissue and ECM dynamics, suggests the existence of a complex, body-wide regulatory mechanism initiated by local tissue loss. In mice, a systemic protease activated by tissue injury relays a signal to stem cells in non-injured tissues to enter an activated state, enhancing their ability to repair damage (51). Post-injury, long-distance communication can also be orchestrated by other factors, including Erk activity (14,52,53), the JAK/STAT pathway (54) and the engagement of the peripheral nervous system (55). However, the discovery of a scaled response in regeneration in our study indicates that the sea anemone *Nematostella* can interpret varying degrees of tissue loss and translate this into scaled gene expression, opening new avenues for investigating the mechanisms of such adaptability. In summary, our findings reveal a complex interplay of local and systemic responses in regeneration, underscoring the need for comprehensive organism-level studies to fully understand the mechanisms of regenerative capabilities across species.

## Acknowledgements

We thank Aliaksandr Halavatyi, Manuel Gunkel, Marko Lampe, Beate Neumann, and Stefan Terjung at the EMBL ALMF for the imaging support. We thank Muzamil Majid Khan for the discussion about the ECM and for sharing ECM-related reagents and the Genomics Core facility at EMBL for RNA sequencing. We also thank Michael Dorrity, Suat Oezbek, Hanh Vu, Jaroslav Ferenc, and Mohannad Dardirry for their comments on the manuscript.

## Author contributions

AI conceptualized the project with SC, DA, TG, PS, and OS’s input. OS and AI supervised the project. SC and PS performed the tomo-seq experiments with HCK’s support. DB and TG performed tomo-seq data analysis. TG generated tomo-seq data visualization. RB established the HCR protocol in Nematostella. SC and PS performed and imaged HCR experiments. MAB performed multiplexed HCRs. SC and PH conducted Edu experiments. SC performed tissue photoconversion experiments. PS established and performed *in vivo* zymography. AP designed the Col4::Dendra2 KI line. SB, PS, and AI established the Col4::Dendra2 KI line. SB performed Col4::Dendra2 photoconversion experiments and quantified the patterns. PS performed all pharmacological experiments. SC, PH, PS, SB, and AI quantified the experimental data. TG designed and assembled all the figures with AI’s support. TG, OS, and AI wrote the manuscript with input from DA. All authors edited the manuscript.

## Materials and Methods

### Animal care and sample collection

Female and male adult animals were kept separately in the dark at 17°C and spawned every 3 weeks as described previously (56). Spawned egg masses were collected and fertilized at room temperature, then kept at 23°C in the dark. Most experiments were done with unfed primary polyps 3 weeks after fertilization.

### Tomography mRNA sequencing

Relaxed polyps were immobilized with 7% MgCl2 in 12 ppt artificial seawater (ASW). Mounting and storage of the samples were done as previously described (57). Changes to the protocol are described below. Polyps were transferred via mouth pipette into the Tissue Freezing Medium (Leica) and oriented using a hair tool so that its tentacles were fully extended parallel to the body column. The ends of the animal were marked with red polyethylene microspheres (Cospheric, REDPMS-0.98 180-212 µm). Samples were frozen in liquid nitrogen and stored at -80°C. Samples were warmed to -20°C prior to cryosectioning into 20-micron thick slices. Each slice was transferred into a separate well of a 96-well tomo-seq plate (Single Cell Discoveries, scdiscoveries.com). Processing of the plates and library preparation was then carried out by Single Cell Discoveries following a CEL-seq2 protocol (58) adapted for a low-input robotics system. Libraries were then multiplexed and sequenced on the Illumina NextSeq500 platform using the 40 bp paired-end setup. A sequencing depth of 50-80 million reads was generated per library.

### *in situ* hybridization chain reaction (HCR)

For each gene target, probe sets for *in situ* HCR v3.0 with split-initiator probes were ordered from Molecular Instruments, Inc (molecularinstruments.com). DNA HCR amplifiers, hybridization, wash, and amplification buffers were also purchased from Molecular Instruments. The staining protocol used for *in situ* HCR was adapted from Molecular Instrument’s protocol for whole-mount zebrafish embryos and larvae, based on Choi and colleagues59. Briefly, polyps were fixed with 4% PFA in PTw (1x PBS, 0.1% tween-20) for 1 hour at room temperature. Samples were then permeabilized with 10% DMSO in PBS, followed by PTx0.5 (1x PBS, 0.5% Triton X-100). Tissue was then clarified via MeOH washes (30%, 60%, 100%) and stored at -20°C for >1 hour before rehydration back into PTw. Samples were then treated with 10 µg/ml Proteinase K (Promega V3021) in PTw for 30min. Excess Proteinase K was removed by washing with PTw. Animals were then refixed in 4% PFA in PTw for 25 min and washed 3 times with PTw. Prehybridization was done in 200 µL Probe Hybridization Buffer (Molecular Instruments) for 30 min at 37°C before hybridizing with the *in situ* probe set (2 pmol in 200 µL Probe hybridization Buffer) overnight at 37°C. In the case of four-channel HCR, Proteinase K treatment and post-fixation were omitted from the protocol. Post-hybridization washes are as follows: 2 times 30 min washes with 400 µL Probe Wash Buffer (Molecular Instruments) at 37°C, then 2 times 5 min washes with 500 µL 5X SSCT (5x SSC pH 7, 0.1% Tween-20) at room temperature. Amplifier hairpins h1 and h2 (18 pmol) were heated separately to 95°C for 90 sec, then snap-cooled to room temperature before being added to 300 µL Probe Amplification Buffer (Molecular Instruments). Samples were then incubated with this amplification solution overnight at room temperature. Excess hairpins were removed by the following washes: 2 times 5 min, then 2 times 30 min with 500 µL 5X SSCT. DNA counterstaining was done by incubating samples overnight with Hoechst (1:1000) in PTw at 4°C. Samples were then mounted into 85% glycerol or Vectashield Plus (Vector Laboratories) or Slowfade Glass (Thermofisher) for imaging. Confocal stacks of whole polyps were acquired on a Leica TCS SP8 confocal microscope using a 40x 1.1NA water objective using the tile scan function, or on a Leica Stellaris 8 using a 20x 0.75NA multi-immersion objective with Type F immersion oil. The acquired z-stack images were processed using Fiji (https://imagej.net/software/fiji/). For quantifying the HCR signal intensity, a sum intensity projection was applied to each image. This method involves summing up the pixel intensities across all slices of the z-stack, providing a cumulative measure of the signal intensity throughout the volume of each polyp. Using the straight line tool, the main body axis of each polyp in the projected image was delineated with a line of 150 µm width. This line served as the basis for extracting the fluorescence intensity profile. The ‘Plot Profile’ function in Fiji was then used to generate a graph representing the HCR signal intensity along the normalized length of the polyp’s body axis.

### Transgenic animals

Homozygous Fgfrb-eGFP adults were crossed with wild-type adults to generate transgenic polyps for staging foot regeneration. Full details on the generation of this transgenic line have been reported previously (60).

### CRISPR/Cas9 mediated knock-in

The Dendra2::3xFlag::col4a2 line was generated using CRISPR/Cas918 using sgRNA AP82 (spacer targeted: gcgtgtgttcccagatgtat) and PCR donor generated with primers AP677/678 (5’Biotin::taagtgccaaggctgccaggcgtgtgttcccagaaacaccccaggaatcaacc and 5’Biotin::gtttttactcacccgatcgcctttctcaccaatacacttgtcatcgtcatccttg) on plasmid AP635 (Addgene #99485). Genotyping shows scarless insertion in the coding sequence, but a partial repeat of the left homology arm was detected in the upstream intron.

### Sample fixation and immunohistochemistry

Animals were fixed and stained using a modified protocol adapted from Genikhovich and Technau61. Briefly, samples were fixed in a solution of 4% paraformaldehyde (PFA; Electron Microscopy Sciences E15710) in Ptw (1x PBS, 0.1% tween-20) for 1 hour at room temperature before being permeabilized with 10% DMSO in PBS for 20 min followed by Ptx 0.5 (1X PBS, 0.5% TritonX-100) for 20 min. Then they were incubated in blocking buffer (5% normal goat serum, 1% BSA, 1% DMSO, 0.1% TritonX-100 in 1X PBS) for 1 hour at room temperature before incubating with the primary antibody (anti-eGFP 1:500, Torry Pines TP401) overnight at 4°C. Samples were then washed 3 times with PTw before incubating with the secondary antibody (goat anti-rabbit alexa488, 1:500, Thermo Fisher A-11008) and DNA stain (Hoechst34580 1:1000 Sigma Aldrich 63493). Samples were then washed with PTw 3 times and mounted into 85% glycerol or Vectashield Plus for imaging.

### EdU staining and quantification

Staining with EdU was done as described previously (60). Briefly, animals were incubated with 50 µM EdU in 12 ppt (parts per thousand) ASW (artificial sea water; seasalt from Instant Ocean) for 30 minutes before immobilizing with 7% MgCl2 solution. Animals were then fixed with 4% PFA in PTw for 1 hour at room temperature before being permeabilized with 10% DMSO in PBS for 20 min followed by Ptx 0.5 (1X PBS, 0.5% TritonX-100) for 20 min. Samples were then blocked for 15 min using 5% BSA (Sigma A2153-50G) in PBS. The Click-it reaction was then performed with alexa647-azide with the sample as specified in the Click-it EdU Cell Proliferation Kit for Imaging (Thermo Fisher C10340) for 30 min at room temperature. Excess labelling reagents were washed away with PTw for 5 min before mounting into 85% glycerol for imaging. EdU+ cells were quantified using FIJI. First a difference of gaussians was calculated for each sample by subtracting the gaussian blurred (sigma = 2) maximum projection image from the gaussian blurred (sigma = 4) maximum projection image. The “find maxima” function was then used to detect individual EdU+ cells and determine their location (x, y coordinates) along the animal body. The 2D Coordinates were then collapsed into 1D by taking only the coordinates corresponding to the oral-aboral axis. The coordinates were then normalized to the size of the animal for each sample to generate a uniform coordinate axis. The numbers of EdU+ cells were then binned into 100 bins and plotted as a heat map using ggplot2.

### Tomo-seq data processing

Reads were mapped to the Nematostella vectensis transcriptome62 using the kallisto-BUStools method63. Full gene sequences were used for mapping as it resulted in consistently higher mapping rates compared to spliced transcripts when using exon-intron boundaries available in the genome annotation. Barcode (8bp) and UMI (6bp) locations in the sequence of the first read were specified accordingly for pseudoalignment. Barcodes were corrected to account for 1 substitution using the white list of barcodes. UMI were then counted for each gene to provide a count matrix for each sample.

### Quality control & sample boundaries

Tomo-seq data for each polyp includes gene counts for 96 slices, some of which do not contain the polyp due to different alignment of polyps for the cutting process and varying polyp sizes. In addition, differences in tissue quality and sequencing depth lead to varying numbers of transcript counts detected per slice (Table 1) which made a dynamic threshold selection for each polyp needed. Sample boundaries were thus defined automatically using a dynamic threshold for each polyp defined as a local minimum of the density function of the number of genes per slice. Slices with high background signals above this threshold were marked and consequently filtered out manually (see Fig. 1a). A summary statistic of the remaining slices is shown in Table 1.

### Anatomical marker genes selection in the reference animal

Gene counts for the genes detected in at least three animals were normalized by total expression per slice and log-transformed. The top 500 highly variable genes (HVGs) as identified for the reference polyp were considered as anatomical markers (Polyp 3 - HH7T7BGXF-42, see Fig. S1c). HGVs were identified by using the FindVariableFeatures function with vst selection method as implemented in ‘Seurat’ v4 (64).

### Reference-based polyp alignment using dynamic time warping

Polyp alignment using dynamic time warping was conducted using the implementation in the DTW package v1.2265. In particular, an open-ended alignment with an asymmetric step function was performed. Thus, we allowed for the alignment of multiple slices within the query polyp to any given slice within the reference polyp (Polyp 3 - b2-42, see Fig. S1c, Table 1) in order to account for regions along the polyps that were longer or shorter than in the reference polyp (Table 1).

### Reference-free polyp alignment using Harmony integration

The R package ‘Seurat’ (v4) (64) was mostly used for analyses and visualizations of tomo-seq data unless stated differently. All slice’ transcriptomes were combined in a Seurat object. A Principle Component Analysis (PCA) was performed and the first 30 PCs were used for removing batch effects using Harmony26. The first 30 Harmony components, excluding the first component (only encoding for batch differences), were used as input for Louvain clustering (resolution of 0.3) and dimensionality reduction by a UMAP embedding. For the analysis of homeostatic polyps, the 6 uncut samples were extracted from the object, and the first 20 Harmony components, excluding the first component (only encoding for batch differences), were used as input for Louvain clustering (resolution of 0.8) and dimensionality reduction by a UMAP embedding.

### Generating SPRING embeddings, in silico polyps and assigning pseudo-locations

The same Harmony components as for the UMAP embeddings were used for generating the SPRING embeddings by using the SPRING web browser tool with no further filtering and by using 5 PC dimensions and 5 nearest neighbors for graph construction and applying the dynamic mode (https://kleintools.hms.harvard.edu/tools/spring.html)27. The R package ‘destiny’ (28) was employed to estimate pseudo-locations by making use of the capacity to order cells or, as here, slices on a gradual trajectory. Cluster identities were used to guide the pseudo-location estimation and slices were finally ranked. Outlines of the in silico polyps were manually generated and combined with the respective heatmap presenting gene expression values ordered by pseudo-location.

### Finding additional spatial markers with the reference-free aligned Tomo-seq data

The FindAllMarkers function was used to identify genes that are positively differentially expressed between clusters of the intact data identified based on a Wilcoxon Rank Sum test (log2FC > 0.3, Fig. 1e, Table 2). By manually inspecting, cluster 1 markers of the uncut data were not identified to be very specific and, therefore, more cluster-specific genes were identified by finding genes that correlate with the top cluster 1 marker NVEC200-000547.1 (Table 2). Correlation instead of cluster 1 markers was used for visualizations in the manuscript. For visualizing genes, either Seurat’s built-in plotting functions were used or the R package Scillus (https://scillus.netlify.app) for generating heatmaps.

### Calculating a cell cycle score

A cell cycle score was calculated for uncut polyps by using Seurat’s AddModuleScore function with default settings after manually intersecting canonical cell cycle markers provided by Seurat with annotated genes in Nematostella by name (see Fig. S4a).

### Quantifying differential systemic expression changes between time points

We performed two analyses to identify differentially expressed genes between time points. i) The first global body-wide differential expression analysis between regeneration time points and the intact state identified 113 differentially expressed (DE) genes for 12 hpa, 122 for 24 hpa, 110 for 48 hpa and 98 for 96 hpa (Wilcoxon Rank Sum test, log2FC > 1) (Table 3), suggesting a lasting effect of injury on gene expression. In addition, time point specific DE genes were calculated by using the FindAllMarkers function in Seurat (Wilcoxon Rank Sum test, log2FC > 0.3; Table 3). In both methods we found a huge overlap of genes between adjacent timepoints and decided to pursue another DE analysis strategy. ii) We searched for adjacent time point markers by applying the FindingMarkers function in Seurat and asking for combined time point specific markers against all other time points (12/24 hpa: early, 24/48 hpa: medium, 48/96 hpa: late) (Wilcoxon Rank Sum test, log2FC > 0.25 & < -0.25; Table 3). Top 50 positive markers for each of the three DE analyses were used as input for a GO term enrichment analysis using the R package ‘GOseq’ (39) (Table 3). The p value cutoff for significant GO terms was 0.05 and the enriched terms belonging to the category ‘biological process’ (BP) were visualized using ‘Revigo’ (66) by automatically lumping similar terms.

### Quantifying differential local expression changes between time points

Log-normalized counts for gene expression for polyps in the common coordinate space obtained by dynamic time warping were used in a GPCounts model (38) to fit Gaussian processes. Briefly, when comparing two-time points, three models are considered: one per time point and one joint model. The log-likelihood of the joint model is then compared to the log-likelihoods of individual models, and a chi-squared test is used to determine the significance of the difference. Significant genes (FDR<1%) are chosen as spatially differentially expressed (DE) between these time points. For each of the spatially DE genes at each time point, the distance between the mean values of the fit and the baseline (the mean of the fit in uncut samples) was then calculated as follows: d(gi,tp1,gi,tp2) where i refers to equally spaced numbers in the (0, 1) interval along the animal (normalized position value), tp1 and tp2 denote two time points and d is a distance metric (Euclidean distance). GO term enrichment analysis was performed using the R package ‘GOseq’ (39) with all 702 DE genes that vary in space and time as identified by Gaussian fitting being used as input. The p-value cutoff applied was 0.02.

### Photoconversion of Kaede

Kaede mRNA was produced using the HiScribe™ T7 ARCA mRNA Kit (with tailing) by New England Biolabs (Catalog #E2060S) from a PCR-amplified template of the Kaede-NLS plasmid sourced from Addgene (Plasmid #57319). After mRNA synthesis, it was purified using SPRISelect magnetic beads from Beckman Coulter (Catalog #B23319). Subsequently, a solution containing the synthesized Kaede-NLS mRNA at a concentration of 200 ng/µl and fluorescein isothiocyanate (FITC) from Thermo Fisher (Catalog #46425) was injected into fertilized eggs. For the photoconversion of Kaede, anesthetized polyps were placed under a Zeiss LSM 780 microscope. Using the microscope’s Plan-Apochromat 20x/0.8 objective, the ‘bleaching’ and ‘regions’ settings were utilized. The conversion was carried out with a 405 nm laser set to 0.6% of its power for 80 iterations. The imaging to observe the results post-photoconversion was done using either a Zeiss LSM 780 or 880 microscope, equipped with the same type of objective lens.

### Quantification of Col4::dendra dynamics

Dendra2, a photoconvertible protein that transforms from green (488 nm) to red (561 nm) upon excitation with UV (405nm), was endogenously tagged to Col 4. Polyps (3 weeks old) maintained at 23°C expressing Col4::Dendra2 were anesthetized with 7% MgCl2 in 12 ppt ASW and mounted on 35 mm dishes (MatTek, P35G-1.5-14-C). Col4::Dendra2 was photoconverted at 4 different regions along the oral-aboral axis using a Zeiss LSM880 Airy confocal microscope. Following that, the animals were washed and returned to 12 ppt ASW. The aboral pole amputation was done by slicing off tissue below the mesenteries with a sharp knife. The polyps were anesthetized 18 hours post amputation (HPA) and imaged with the same microscope. To understand the dynamics of Col4, the positions of the photoconverted clones (xk) were recorded for the control and regenerating polyps after photoconversion (t = t0) and 18 hours post amputation (HPA) (t = t1). The length of every polyp was normalized from oral pole (x = 0) to aboral pole (x = 1) with the length of the animals at t0. This allows quantifying a relative velocity field for Col4::Dendra2 by computing v(xk,t) = xk(t0) - xk(t1), where k denotes the position of the clone along the axis. The plot approximates a field of Col4 remodeling during regeneration, with the x-axis representing axial position and y-axis representing the velocity of Col4::Dendra2 towards oral (positive) or aboral pole (negative). The shaded area represents a 95% confidence interval with the fitting of a linear model.

### *In vivo* zymography

Intact, foot-amputated, and mid-body amputated polyps were incubated in 12 ppt ASW in the well of a 48-well plate with the DQ-Collagen 4, FITC conjugate substrate (Thermo Fisher #D12052; final concentration 20 µg/mL) in the dark for 18 hours at room temperature. After removing the substrate solution, two washing steps were performed in the dark with 12 ppt ASW for 5 min. Then, the animals were anesthetized with 7% MgCl2 and fixed for 30 min with ice-cold 4% paraformaldehyde/PBS at room temperature and washed 2x 5 min with PBS. The polyps were kept for at least o/n at 4°C in a Vectashield mounting medium with DAPI (Vector Laboratories) before being mounted and imaged on a Leica SP8 confocal system. Intensities were measured in the total area of the body column using an equal number of z-projected stacks for each animal. Image analysis was performed with FIJI.

### Inhibitors of matrix metalloproteases

Following spawning, Fgfrb-eGFP embryos were grown at 23°C. At 10 dpf, primary polyps were transferred to a 3cm plastic petri dish containing 15 mL of 0.05% DMSO in 12 ppt ASW or 15 mL of 50 µM GM6001 in 12 ppt ASW. After a 30 min treatment at room temperature, the heads of the polyps were removed with a cut below the pharynx using a sharp knife. The heads and any non-cut polyps or polyps with misplaced cuts, were removed from the dish. All other polyps were allowed to recover at 23°C with daily replacement of the solutions. At 5 dpa (=120 hpa), polyps were immobilized by carefully adding 7% MgCl2 to the dish and fixed for 1h at room temperature in 4% PFA in 12 ppt ASW. Samples were then permeabilized in 10% DMSO and PTx 0.2%, before blocking a specific antibody staining using a blocking buffer containing 1% BSA and 5% goat serum. Fgfrb-eGFP signal was detected using an anti-GFP antibody (TP401, Torrey Pine Biolabs) in combination with nuclear (Hoechst) and F-actin staining (phalloidin). Images were taken on a Leica SP8-CSU confocal microscope.

### Data availability

Raw sequencing data is deposited on NCBI SRA under the BioProject accession code PRJNA1079674 and available upon request. The R code and the processed data used to analyze the data and to generate plots presented is publicly available on github (https://github.com/tobiasgerber/NematostellaTomo).

## Supplementary Figures

**Fig. S1.**
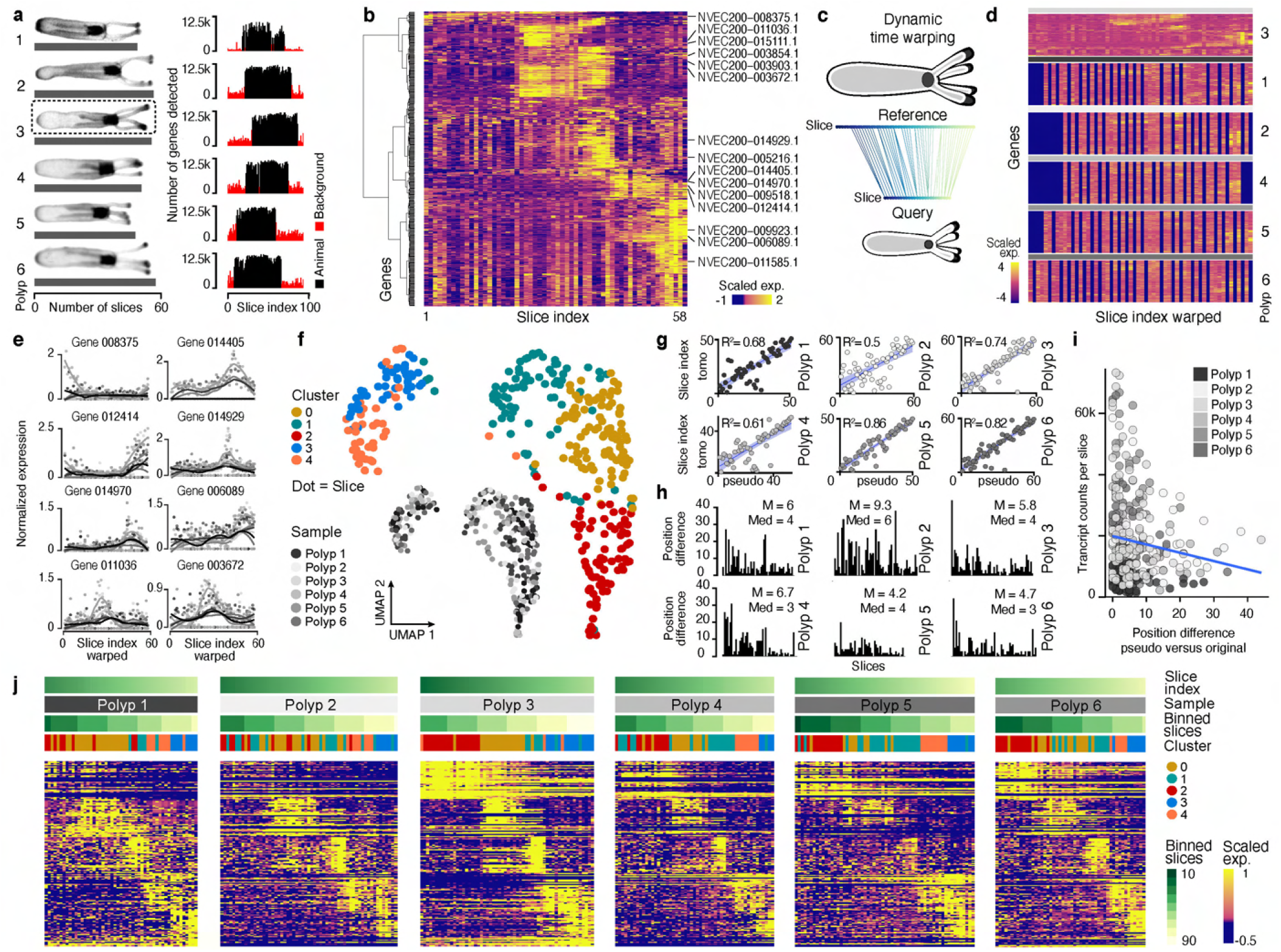
Analyses of tomo-seq data using both reference-based and unbiased ordering of slices. (**a**) Brightfield images of the 6 intact polyps studied (left). Up to 60 high-quality slices are obtained from each polyp. Right: Histogram indicates the number of genes detected across all the slices obtained. Black bars denote slices that included polyps, which were identified by higher RNA counts. (**b**) Brightfield images of the 6 intact polyps studied (left). Up to 60 high-quality slices are obtained from each polyp. Right: Histogram indicates the number of genes detected across all the slices obtained. Black bars denote slices that included polyps, which were identified by higher RNA counts. (**c**) MSchematic explaining the principle of the dynamic time warping approach employed for reference-guided integration. The sliced index of a query animal is mapped (warped) to match a selected reference animal. (**d**) Genes in b visualized as heatmaps for time-warped slices across all polyps. Slices are aligned in a combined coordinate system based on the reference polyp 3. Dark blue slices indicate non-matching coordinates in the query polyps. (**e**) Gene expression patterns of representative genes visualized as a function of the warped slice index for all six polyps. Points correspond to normalized expression counts; solid lines denote loess-smoothed curves. (**f**) UMAP embedding of the RNA profiles of individual slices after Harmony-based integration across all polyps. Colors denote the identity of Louvain clusters. Inset shows the same embedding with color denoting the source by polyp identity. (**g**) Comparison of the pseudo temporal ordering estimated based on HARMONY aligned data to the observed tomo-seq slice indexes for individual polyps. Solid lines represent linear regression fits with R2 values shown. (**h**) Histograms of the absolute positional divergence between (in slices) the observed tomo-seq and estimated pseudo temporal orderings for the individual polyp. Mean and median values are provided. (**i**) Scatter plot between the positional divergence is in h versus the number of detected transcripts per slice. Color denotes polyp. A solid blue line denotes linear regression fit. (**j**) Cluster-specific marker genes, as in Fig. 1g, are presented as heatmaps for each polyp individually, with slices being ordered by the original slice index.

**Fig. S2.**
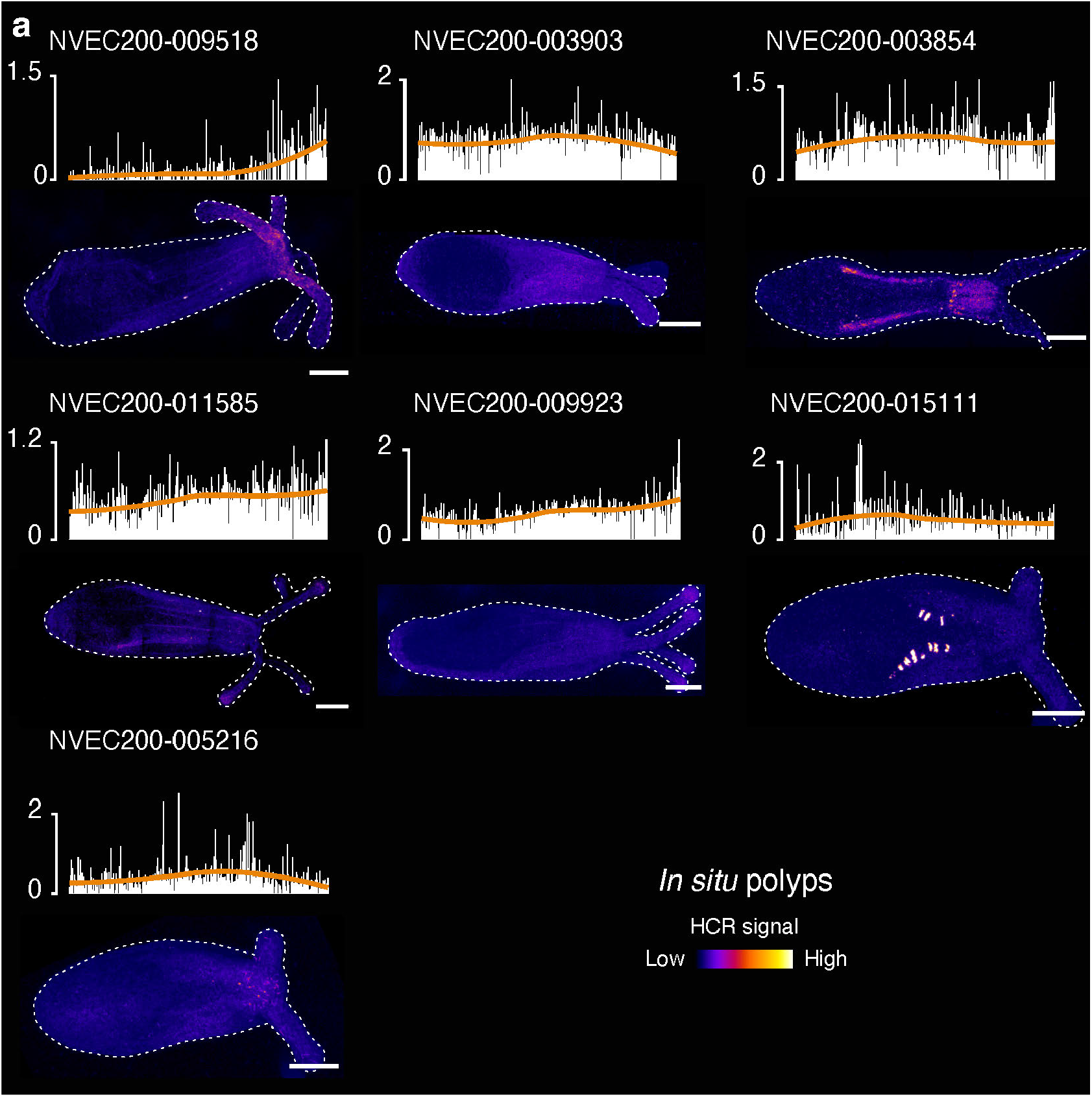
Validation of tomo-seq data with HCR and representation of expression patterns for select genes using in silico polyps. (**a**) Histograms of normalized gene expressions per slice ordered by pseudo-location (top rows) and maximum projection images of whole mount HCR *in situ* (bottom rows) for additional genes. HCR intensities are converted by pseudocolor palette ‘fire’. Scale bars: 100 µm.

**Fig. S3.**
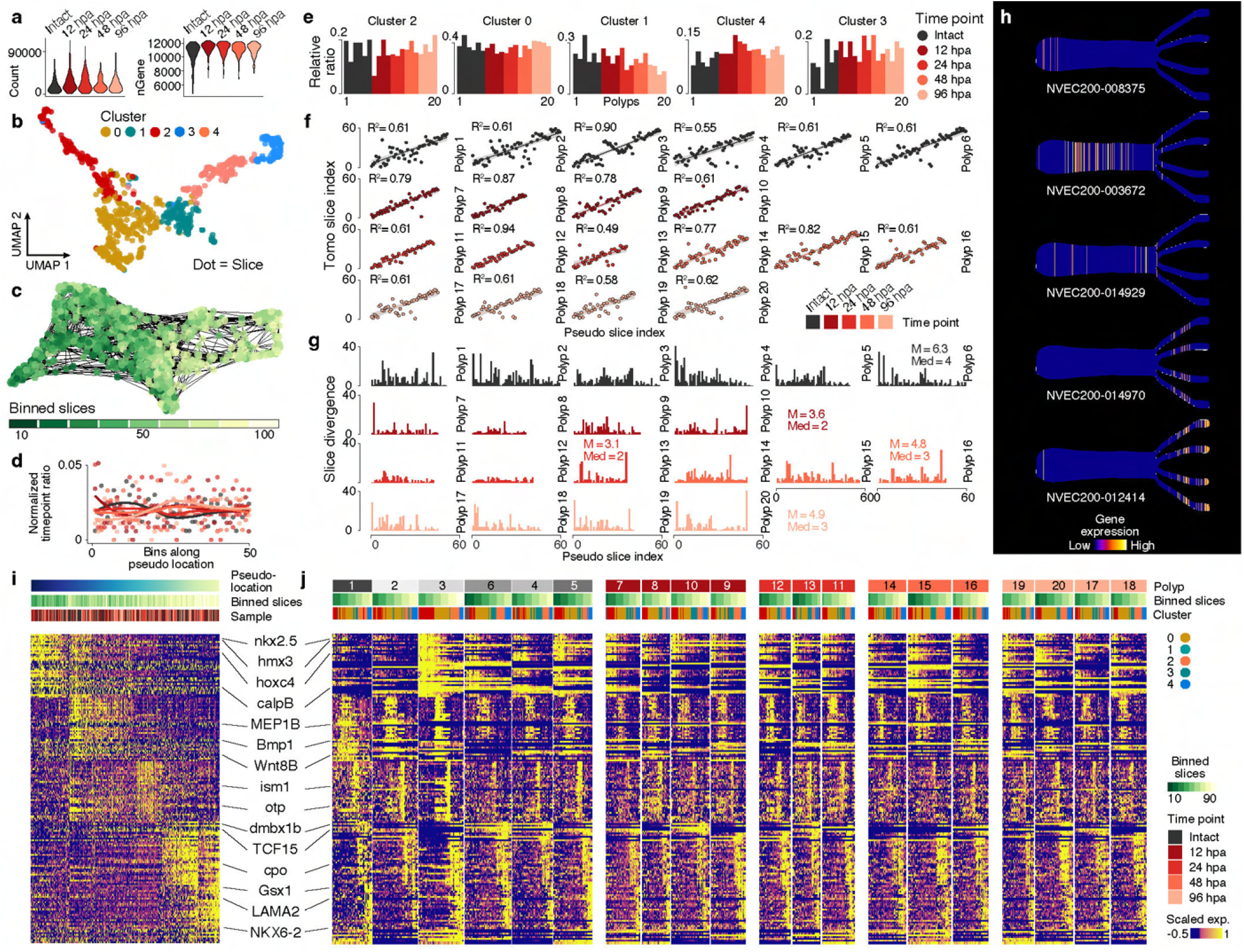
Molecular landscape of foot-regenerating polyps. (**a**) Violin plots showing the number of counts (left) and genes (right) detected per slice. (**b**) UMAP embedding of slices after data integration of 957 tomo-seq slices across all time points (0 - 96 hpa). Colors represent Louvain cluster identity. (**c**) SPRING embedding of tomo-seq slices colored by their original slice index bin. Grey lines show the connection of slice transcriptomes in the SPRING k-nearest-neighbor graph. (**d**) Slices are binned along the pseudo-locations, and the normalized time point contribution per bin is visualized as a scatter plot with a loess fit denoted as a line per time point. (**e**) Histograms represent the relative time point contribution per polyp to the clusters in b. (**f**) Scatter plots compare pseudo rank ordering to actual tomo-seq slice indexes with slices being ranked within each polyp. (**g**) Histograms show the position divergence between the two positions per slice. Mean (M) and median (Med) values are provided. (**h**) In silico polyps visualize site-specific expression patterns as revealed by HCR staining and expression histograms of respective genes as shown in Fig. 1f. (**i**) Heatmap representation of gene expression for top cluster specific marker genes identified for the intact polyps across all slices (see Fig. 1e). Slices are ordered by pseudo-location as presented in Fig. 2b. (**j**) The same genes as in g are plotted as heatmaps for each polyp individually with slices being ordered by original slice index.

**Fig. S4.**
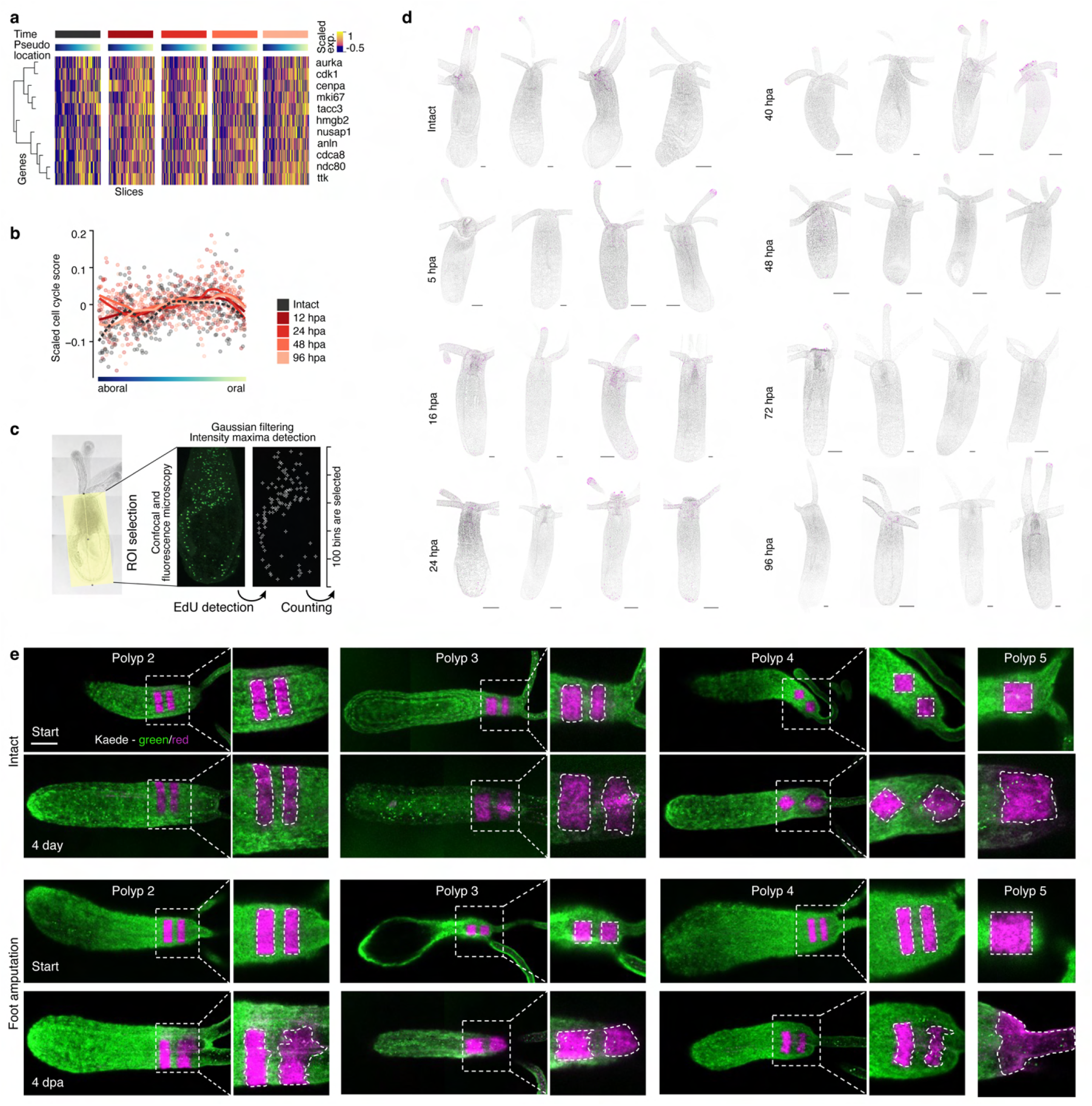
Epithelial dynamics of foot-regenerating polyps. (**a**) Heatmap showing the scaled gene expression levels of canonical cell cycle genes (rows) for each slice (columns) that were used to calculate a cell cycle score in b. Slices are separated by time point and ordered by pseudo-location ranking (see Fig. 2b). Hierarchical clustering of genes is shown as a dendrogram. (**b**) Genes in panel a were used to calculate a cell cycle score, which is visualized as a scatter plot ordered by pseudo-location. Lines represent fitted loess curves to smooth the scattered data. (**c**) Image analysis workflow of maximum projection of EdU-positive cells in polyps. (**d**) EdU/Hoechst maximum projection images for intact and foot-regenerating polyps at the indicated time points. Four polyps stained with Hoechst (gray) and EdU (magenta) are analyzed per time point during regeneration. (**e**) Maximum projection images of photo-converted oral tissue patches in additional intact (top) and foot-regenerating (bottom) polyps expressing Kaede (green: not converted; magenta: photo-converted). The pattern of the photo-converted patches is highlighted in dashed lines and shown shortly after the photoconversion (start) and four days post-conversion. Scale bars: 100 µm.

**Fig. S5.**
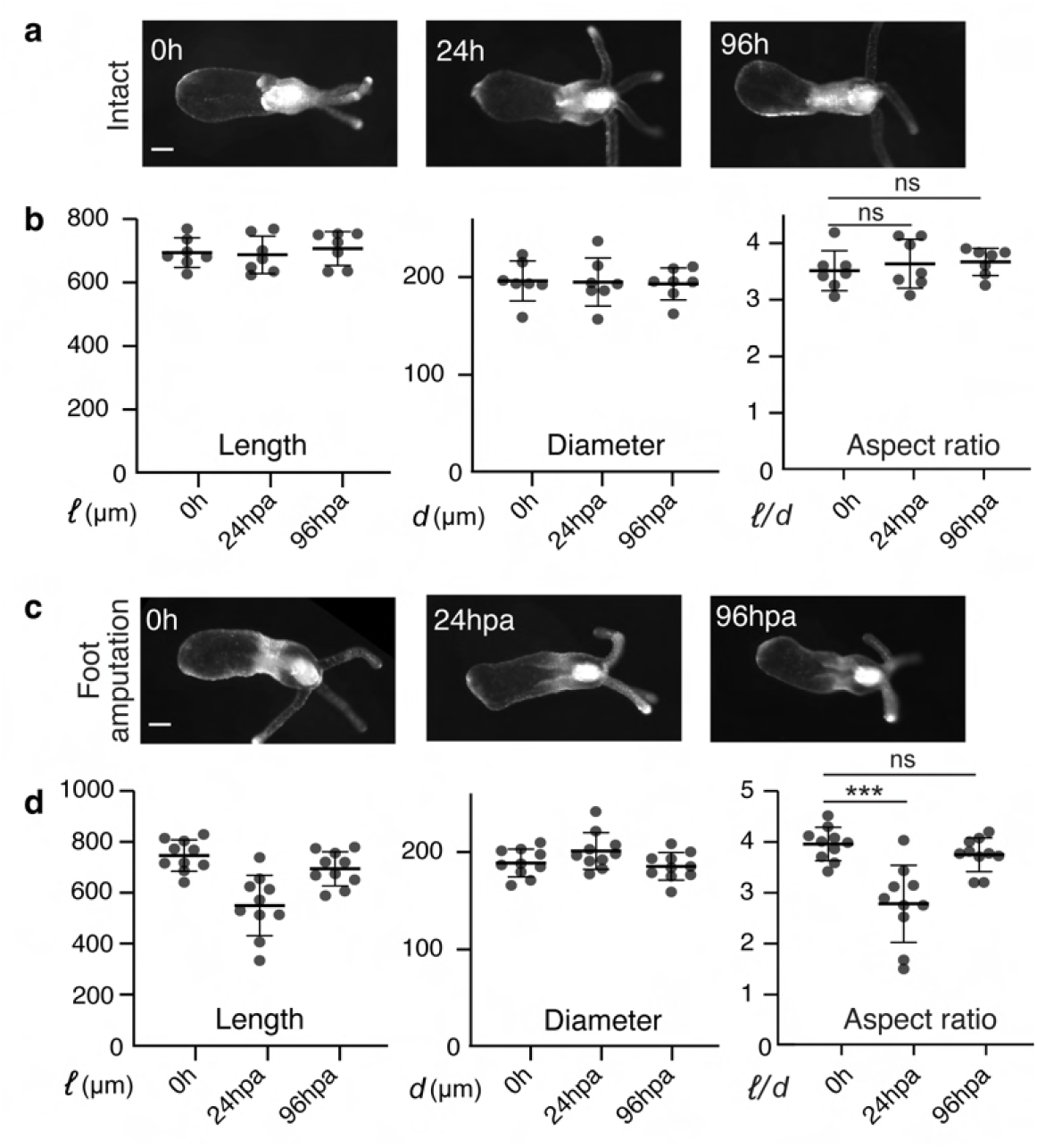
Foot-regenerating polyps re-adjust their aspect ratio. (**a**) Brightfield images show the same intact polyps over the indicated time points. Scale bar: 100 µm. (**b**) Scatter plots showing Length, diameter, and aspect ratios in intact polyps (n= 7 polyps). ns = not significant. (**c**) Brightfield images show the same foot-regenerating polyp at each indicated time point. (**d**) Scatter plots show length, diameter, and aspect ratios in foot-regenerating polyps (n= 10 polyps). Data in b and d are shown as individual values (Mean with SD; unpaired Student’s two-tailed t-test, ^***^p <0.003) Scale bars: 100 µm.

**Fig. S6.**
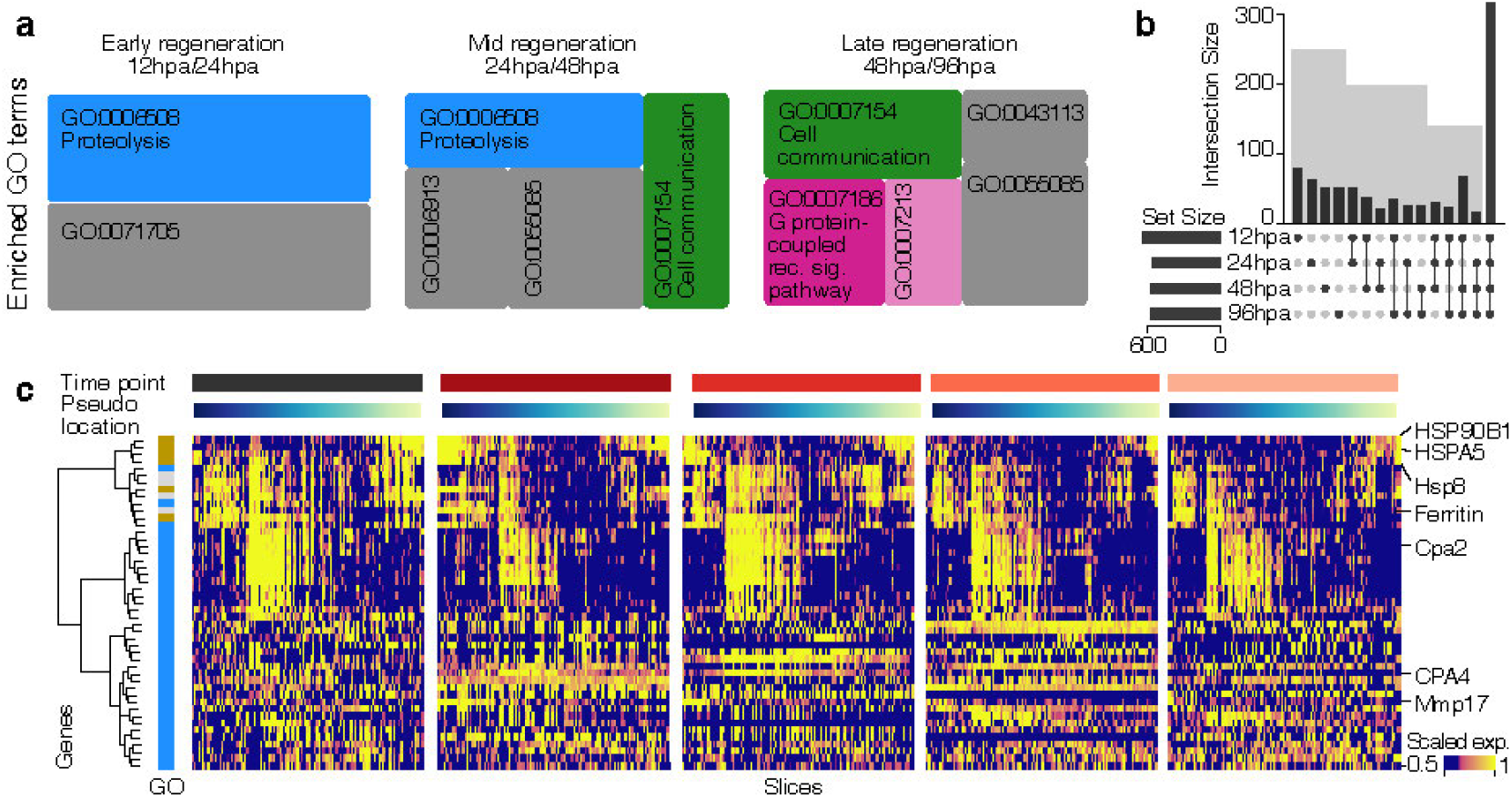
Enriched GO terms of biological processes in tomo-seq data of foot-regenerating polyps. (**a**) Enriched GO terms of biological processes (BP) for sets of genes associated with evidence for upregulation during distinct regenerative stages. Enrichment analysis was conducted using GOseq (39), considering positively differentially expressed genes between the intact state and different regeneration time points (always combining pairs of time points; 12/24 hpa, mid 24/48 hpa, and late 48/96 hpa; Table 2). Shade variations of the same color indicate that terms belong to the same term category. (**b**) Overlap of genes with evidence for positionally local differential gene expression (Fig. 3e), comparing spatial gene expression for each of 4 time points post-amputation with the intact expression pattern. Shown is an upset plot, illustrating overlapping DE genes (out of a total of N=702 spatially differentially expressed genes, FDR<1%). (**c**)Visualization of spatial expression patterns for genes that belong to the top enriched GO terms in Fig. 3f, displayed as heatmaps across time points ordered by pseudo-location, respectively. The colored sidebar refers to GO terms in Fig. 3f.

**Fig. S7.**
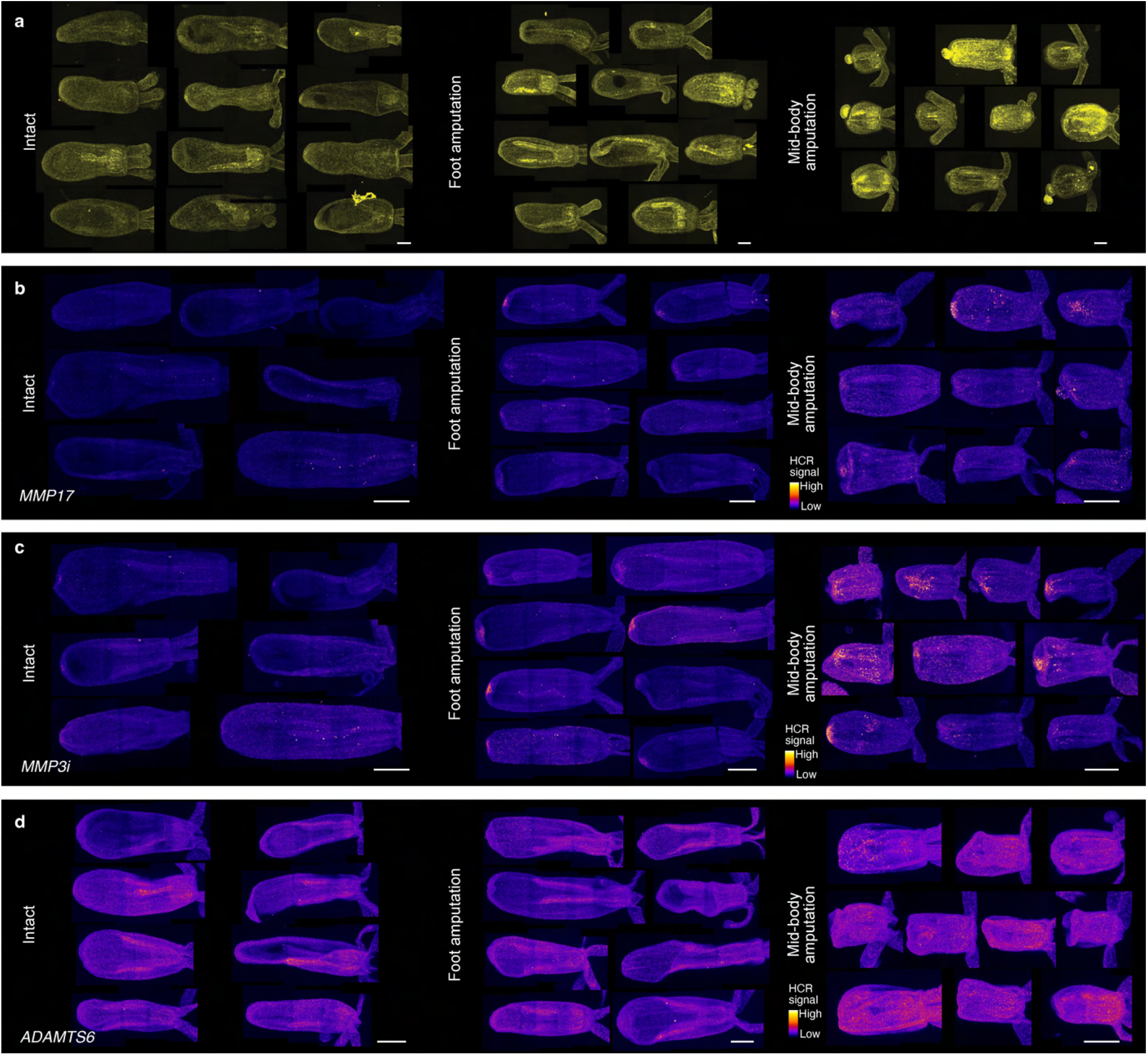
Expression and activity of metalloproteases. (**a**) Maximum projection images of intact (control), foot-amputated, and mid-body amputated polyps incubated with DQ-Col 4 for *in vivo* zymography. Intensities were used to generate the diagram in Fig. 4a. Scale bar: 100 µm. (**b-d**) Maximum projection images of whole mount HCR *in situ* for MMP17, MMP3i, and ADAMTS6 before (left) and after foot amputation (center) and after mid-body amputation (right). The intensity profile for each polyp was used as input for the plot in Fig. 4d. HCR intensity profiles are shown as pseudocolor palette ‘fire’. Scale bars: 100 µm.

**Fig. S8.**
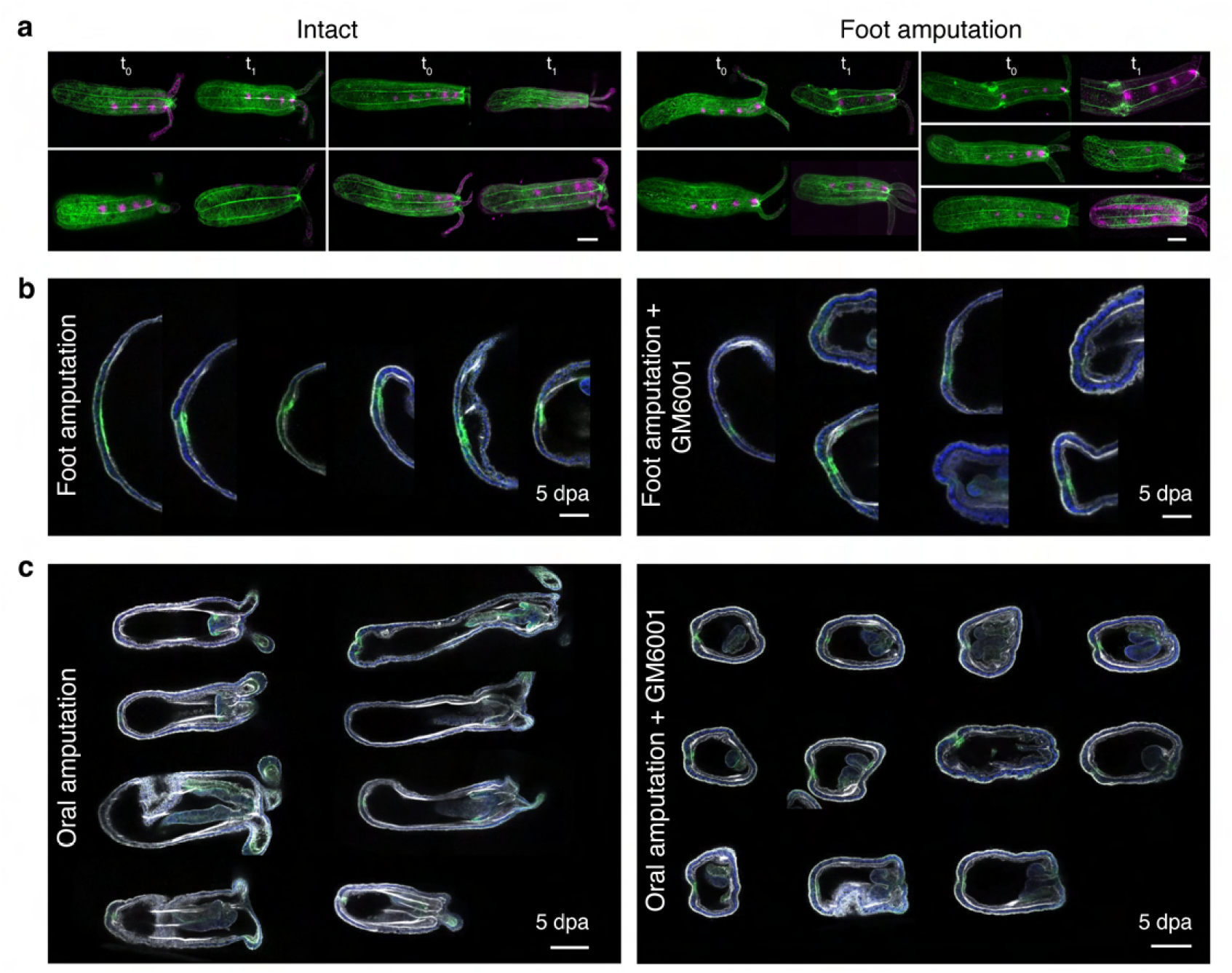
Inhibition of matrix metalloproteases disrupts regeneration. (**a**) Maximum projection images of additional polyps showing Col4::Dendra2 (green) and photoconverted Col4::Dendra2 patches (magenta) along the oral-aboral axis in intact (left) and foot regenerating animals (right). t0: 0 hour, t1: 18 hours. Scale bars: 100 µm. (**b**) Maximum projection images of control (left) and GM6001-treated Fgfrb-eGFP polyps (right) after foot amputation at 5 dpa. Scale bar: 50 µm. (**c**) Maximum projection images of control (left) and GM6001-treated (right) Fgfrb-eGFP polyps after oral amputation. Scale bar: 100 µm.

## References

1 Sun, F. & Poss, K. D. Inter-organ communication during tissue regeneration. Development 150, doi:10.1242/dev.202166 (2023).

2 Losner, J., Courtemanche, K. & Whited, J. L. A cross-species analysis of systemic mediators of repair and complex tissue regeneration. NPJ Regen Med 6, 21, doi:10.1038/s41536-021-00130-6 (2021).

3 Brockes, J. P. & Kumar, A. Comparative aspects of animal regeneration. Annu Rev Cell Dev Biol 24, 525–549, doi:10.1146/annurev.cellbio.24.110707.175336 (2008).

4 Bely, A. E. & Nyberg, K. G. Evolution of animal regeneration: re-emergence of a field. Trends Ecol Evol 25, 161–170, doi:10.1016/j.tree.2009.08.005 (2010).

5 Sanchez Alvarado, A. & Tsonis, P. A. Bridging the regeneration gap: genetic insights from diverse animal models. Nat Rev Genet 7, 873–884, doi:10.1038/nrg1923 (2006).

6 Tanaka, E. M. & Reddien, P. W. The cellular basis for animal regeneration. Dev Cell 21, 172–185, doi:10.1016/j.devcel.2011.06.016 (2011).

7 Poss, K. D. Advances in understanding tissue regenerative capacity and mechanisms in animals. Nat Rev Genet 11, 710–722, doi:10.1038/nrg2879 (2010).

8 Wenemoser, D. & Reddien, P. W. Planarian regeneration involves distinct stem cell responses to wounds and tissue absence. Dev Biol 344, 979–991, doi:10.1016/j.ydbio.2010.06.017 (2010).

9 Sun, F. et al. Enhancer selection dictates gene expression responses in remote organs during tissue regeneration. Nat Cell Biol 24, 685–696, doi:10.1038/s41556-022-00906-y (2022).

10 Rodgers, J. T. et al. mTORC1 controls the adaptive transition of quiescent stem cells from G0 to G(Alert). Nature 510, 393–396, doi:10.1038/nature13255 (2014).

11 Johnson, K., Bateman, J., DiTommaso, T., Wong, A. Y. & Whited, J. L. Systemic cell cycle activation is induced following complex tissue injury in axolotl. Dev Biol 433, 461–472, doi:10.1016/j.ydbio.2017.07.010 (2018).

12 DuBuc, T. Q., Traylor-Knowles, N. & Martindale, M. Q. Initiating a regenerative response; cellular and molecular features of wound healing in the cnidarian Nematostella vectensis. BMC Biol 12, 24, doi:10.1186/1741-7007-12-24 (2014).

13 Srivastava, M., Mazza-Curll, K. L., van Wolfswinkel, J. C. & Reddien, P. W. Whole-body acoel regeneration is controlled by Wnt and Bmp-Admp signaling. Curr Biol 24, 1107–1113, doi:10.1016/j.cub.2014.03.042 (2014).

14 Fan, Y. et al. Ultrafast and long-range coordination of wound responses is essential for whole-body regeneration. bioRxiv, doi:10.1101/2023.03.15.532844 (2023).

15 Passamaneck, Y. J. & Martindale, M. Q. Cell proliferation is necessary for the regeneration of oral structures in the anthozoan cnidarian Nematostella vectensis. BMC Dev Biol 12, 34, doi:10.1186/1471-213X-12-34 (2012).

16 Amiel, A. R. et al. Characterization of Morphological and Cellular Events Underlying Oral Regeneration in the Sea Anemone, Nematostella vectensis. Int J Mol Sci 16, 28449–28471, doi:10.3390/ijms161226100 (2015).

17 Ikmi, A., McKinney, S. A., Delventhal, K. M. & Gibson, M. C. TALEN and CRISPR/Cas9-mediated genome editing in the early-branching metazoan Nematostella vectensis. Nat Commun 5, 5486, doi:10.1038/ncomms6486 (2014).

18 Paix, A. et al. Endogenous tagging of multiple cellular components in the sea anemone Nematostella vectensis. Proc Natl Acad Sci U S A 120, e2215958120, doi:10.1073/pnas.2215958120 (2023).

19 Sebe-Pedros, A. et al. Cnidarian Cell Type Diversity and Regulation Revealed by Whole-Organism Single-Cell RNA-Seq. Cell 173, 1520–1534 e1520, doi:10.1016/j.cell.2018.05.019 (2018).

20 Steger, J. et al. Single-cell transcriptomics identifies conserved regulators of neuroglandular lineages. Cell Rep 40, 111370, doi:10.1016/j.celrep.2022.111370 (2022).

21 Hereroa, J. et al. Whole body regeneration deploys a rewired embryonic gene regulatory network logic. bioRxiv, 658930, doi:10.1101/658930 (2021).

22 Kruse, F., Junker, J. P., van Oudenaarden, A. & Bakkers, J. Tomo-seq: A method to obtain genome-wide expression data with spatial resolution. Methods Cell Biol 135, 299–307, doi:10.1016/bs.mcb.2016.01.006 (2016).

23 Junker, J. P. et al. Genome-wide RNA Tomography in the zebrafish embryo. Cell 159, 662–675, doi:10.1016/j.cell.2014.09.038 (2014).

24 Schild, E. S. et al. Spatial transcriptomics of the nematode Caenorhabditis elegans using RNA tomography. STAR Protoc 2, 100411, doi:10.1016/j.xpro.2021.100411 (2021).

25 Giorgino, T. Computing and Visualizing Dynamic Time Warping Alignments in R: The dtw Package. J Stat Softw 31, 1 – 24, doi:10.18637/jss.v031.i07 (2009).

26 Korsunsky, I. et al. Fast, sensitive and accurate integration of single-cell data with Harmony. Nat Methods 16, 1289–1296, doi:10.1038/s41592-019-0619-0 (2019).

27 Weinreb, C., Wolock, S. & Klein, A. M. SPRING: a kinetic interface for visualizing high dimensional single-cell expression data. Bioinformatics 34, 1246–1248, doi:10.1093/bioinformatics/btx792 (2018).

28 Angerer, P. et al. destiny: diffusion maps for large-scale single-cell data in R. Bioinformatics 32, 1241–1243, doi:10.1093/bioinformatics/btv715 (2016).

29 Pukhlyakova, E. A., Kirillova, A. O., Kraus, Y. A., Zimmermann, B. & Technau, U. A cadherin switch marks germ layer formation in the diploblastic sea anemone Nematostella vectensis. Development 146, doi:10.1242/dev.174623 (2019).

30 Mazza, M. E., Pang, K., Martindale, M. Q. & Finnerty, J. R. Genomic organization, gene structure, and developmental expression of three clustered otx genes in the sea anemone Nematostella vectensis. J Exp Zool B Mol Dev Evol 308, 494–506, doi:10.1002/jez.b.21158 (2007).

31 Leclere, L. & Rentzsch, F. RGM regulates BMP-mediated secondary axis formation in the sea anemone Nematostella vectensis. Cell Rep 9, 1921–1930, doi:10.1016/j.celrep.2014.11.009 (2014).

32 Faltine-Gonzalez, D. Z. & Layden, M. J. Characterization of nAChRs in Nematostella vectensis supports neuronal and non-neuronal roles in the cnidarian-bilaterian common ancestor. Evodevo 10, 27, doi:10.1186/s13227-019-0136-3 (2019).

33 Aldine, R. A., Kevin, F., Solène, F. & Eric, R. Synergic coordination of stem cells is required to induce a regenerative response in anthozoan cnidarians. bioRxiv, 2019.2012.2031.891804, doi:10.1101/2019.12.31.891804 (2019).

34 Bradshaw, B., Thompson, K. & Frank, U. Distinct mechanisms underlie oral vs aboral regeneration in the cnidarian Hydractinia echinata. Elife 4, e05506, doi:10.7554/eLife.05506 (2015).

35 Schaffer, A. A., Bazarsky, M., Levy, K., Chalifa-Caspi, V. & Gat, U. A transcriptional time-course analysis of oral vs. aboral whole-body regeneration in the Sea anemone Nematostella vectensis. BMC Genomics 17, 718, doi:10.1186/s12864-016-3027-1 (2016).

36 Kiyozumi, D., Sugimoto, N. & Sekiguchi, K. Breakdown of the reciprocal stabilization of QBRICK/Frem1, Fras1, and Frem2 at the basement membrane provokes Fraser syndrome-like defects. Proc Natl Acad Sci U S A 103, 11981–11986, doi:10.1073/pnas.0601011103 (2006).

37 Jadeja, S. et al. Identification of a new gene mutated in Fraser syndrome and mouse myelencephalic blebs. Nat Genet 37, 520–525, doi:10.1038/ng1549 (2005).

38 BinTayyash, N. et al. Non-parametric modelling of temporal and spatial counts data from RNA-seq experiments. Bioinformatics 37, 3788–3795, doi:10.1093/bioinformatics/btab486 (2021).

39 Young, M. D., Wakefield, M. J., Smyth, G. K. & Oshlack, A. Gene ontology analysis for RNA-seq: accounting for selection bias. Genome Biol 11, R14, doi:10.1186/gb-2010-11-2-r14 (2010).

40 Jeffery, W. R. et al. Differentially expressed chaperone genes reveal a stress response required for unidirectional regeneration in the basal chordate Ciona. BMC Biol 21, 148, doi:10.1186/s12915-023-01633-y (2023).

41 Wang, C. et al. Heat shock protein DNAJA1 stabilizes PIWI proteins to support regeneration and homeostasis of planarian Schmidtea mediterranea. J Biol Chem 294, 9873–9887, doi:10.1074/jbc.RA118.004445 (2019).

42 Pearl, E. J., Barker, D., Day, R. C. & Beck, C. W. Identification of genes associated with regenerative success of Xenopus laevis hindlimbs. BMC Dev Biol 8, 66, doi:10.1186/1471-213X-8-66 (2008).

43 Makino, S. et al. Heat-shock protein 60 is required for blastema formation and maintenance during regeneration. Proc Natl Acad Sci U S A 102, 14599–14604, doi:10.1073/pnas.0507408102 (2005).

44 Vandooren, J., Geurts, N., Martens, E., Van den Steen, P. E. & Opdenakker, G. Zymography methods for visualizing hydrolytic enzymes. Nat Methods 10, 211–220, doi:10.1038/nmeth.2371 (2013).

45 Aufschnaiter, R. et al. In vivo imaging of basement membrane movement: ECM patterning shapes Hydra polyps. J Cell Sci 124, 4027–4038, doi:10.1242/jcs.087239 (2011).

46 Veschgini, M. et al. Wnt/beta-catenin signaling induces axial elasticity patterns of Hydra extracellular matrix. iScience 26, 106416, doi:10.1016/j.isci.2023.106416 (2023).

47 Sarras, M. P., Jr. Components, structure, biogenesis and function of the Hydra extracellular matrix in regeneration, pattern formation and cell differentiation. Int J Dev Biol 56, 567–576, doi:10.1387/ijdb.113445ms (2012).

48 Phan, A. Q. et al. Positional information in axolotl and mouse limb extracellular matrix is mediated via heparan sulfate and fibroblast growth factor during limb regeneration in the axolotl (Ambystoma mexicanum). Regeneration (Oxf) 2, 182–201, doi:10.1002/reg2.40 (2015).

49 Sanchez-Iranzo, H. et al. Transient fibrosis resolves via fibroblast inactivation in the regenerating zebrafish heart. Proc Natl Acad Sci U S A 115, 4188–4193, doi:10.1073/pnas.1716713115 (2018).

50 Sonpho, E. et al. Decellularization Enables Characterization and Functional Analysis of Extracellular Matrix in Planarian Regeneration. Mol Cell Proteomics 20, 100137, doi:10.1016/j.mcpro.2021.100137 (2021).

51 Rodgers, J. T., Schroeder, M. D., Ma, C. & Rando, T. A. HGFA Is an Injury-Regulated Systemic Factor that Induces the Transition of Stem Cells into G(Alert). Cell Rep 19, 479–486, doi:10.1016/j.celrep.2017.03.066 (2017).

52 Hiratsuka, T. et al. Intercellular propagation of extracellular signal-regulated kinase activation revealed by in vivo imaging of mouse skin. eLife 4, e05178, doi:10.7554/eLife.05178 (2015).

53 De Simone, A. et al. Control of osteoblast regeneration by a train of Erk activity waves. Nature 590, 129–133, doi:10.1038/s41586-020-03085-8 (2021).

54 Chakrabarti, S. et al. Remote Control of Intestinal Stem Cell Activity by Haemocytes in Drosophila. PLoS Genet 12, e1006089, doi:10.1371/journal.pgen.1006089 (2016).

55 Duygu, P.-D. et al. Adrenergic signaling stimulates body-wide stem cell activation for limb regeneration. bioRxiv, 2021.2012.2029.474455, doi:10.1101/2021.12.29.474455 (2023).

56 Fritzenwanker, J. H. & Technau, U. Induction of gametogenesis in the basal cnidarian Nematostella vectensis(Anthozoa). Dev Genes Evol 212, 99–103, doi:10.1007/s00427-002-0214-7 (2002).

57 Ebbing, A. et al. Spatial Transcriptomics of C. elegans Males and Hermaphrodites Identifies Sex-Specific Differences in Gene Expression Patterns. Dev Cell 47, 801–813 e806, doi:10.1016/j.devcel.2018.10.016 (2018).

58 Hashimshony, T. et al. CEL-Seq2: sensitive highly-multiplexed single-cell RNA-Seq. Genome Biol 17, 77, doi:10.1186/s13059-016-0938-8 (2016).

59 Choi, H. M. T. et al. Third-generation in situ hybridization chain reaction: multiplexed, quantitative, sensitive, versatile, robust. Development 145, doi:10.1242/dev.165753 (2018).

60 Ikmi, A. et al. Feeding-dependent tentacle development in the sea anemone Nematostella vectensis. Nat Commun 11, 4399, doi:10.1038/s41467-020-18133-0 (2020).

61 Genikhovich, G. & Technau, U. Induction of spawning in the starlet sea anemone Nematostella vectensis, in vitro fertilization of gametes, and dejellying of zygotes. Cold Spring Harb Protoc 2009, pdb prot5281, doi:10.1101/pdb.prot5281 (2009).

62 Zimmermann, B. et al. Sea anemone genomes reveal ancestral metazoan chromosomal macrosynteny. bioRxiv, 2020.2010.2030.359448, doi:10.1101/2020.10.30.359448 (2023).

63 Melsted, P. et al. Modular, efficient and constant-memory single-cell RNA-seq preprocessing. Nat Biotechnol 39, 813–818, doi:10.1038/s41587-021-00870-2 (2021).

64 Hao, Y. et al. Integrated analysis of multimodal single-cell data. Cell 184, 3573–3587 e3529, doi:10.1016/j.cell.2021.04.048 (2021).

65 Giorgino, T. Computing and Visualizing Dynamic Time Warping Alignments in R: The dtw Package. J Stat Softw 31, 1–24, doi:10.18637/jss.v031.i07 (2009).

66 Supek, F., Bosnjak, M., Skunca, N. & Smuc, T. REVIGO summarizes and visualizes long lists of gene ontology terms. PLoS One 6, e21800, doi:10.1371/journal.pone.0021800 (2011).

